# Loss of the Wnt/β-catenin pathway in microglia of the developing brain drives pro-inflammatory activation leading to white matter injury

**DOI:** 10.1101/334359

**Authors:** Juliette Van Steenwinckel, Anne-Laure Schang, Michelle L Krishnan, Vincent Degos, Andrée Delahaye-Duriez, Cindy Bokobza, Franck Verdonk, Amélie Montané, Stéphanie Sigaut, Olivier Hennebert, Sophie Lebon, Leslie Schwendimann, Tifenn Le Charpentier, Rahma Hassan-Abdi, Gareth Ball, Paul Aljabar, Alka Saxena, Rebecca K Holloway, Walter Birchmeier, Veronique Miron, David Rowitch, Fabrice Chretien, Claire Leconte, Valérie C Besson, Enrico G Petretto, A David Edwards, Henrik Hagberg, Nadia Soussi-Yanicostas, Bobbi Fleiss, Pierre Gressens

## Abstract

Microglia of the developing brain have unique functional properties but how their activation states is regulated is poorly understood. Inflammatory activation of microglia in the still-developing brain of preterm born infants is associated with permanent neurological sequelae in 9 million infants every year. Investigating the regulators of microglial activation in the developing brain with multiple models of neuroinflammation-mediated injury and primary human microglia we found that a reduction in Wnt/β-catenin signalling is necessary and sufficient to drive an oligodendrocyte-injurious microglial phenotype. We validated in a cohort of preterm born infants that genomic variation in the *WNT* pathway is associated with the levels of connectivity found in their brains. Using a Wnt agonist delivered by a BBB penetrant microglia-specific targeting nanocarrier we prevented in our animal model the pro-inflammatory microglial activation, white matter injury and behavioural deficits. Collectively, these data validate that the Wnt pathway regulates microglial activation, is critical in the evolution of an important form of human brain injury and is a viable therapeutic target.

## Introduction

Neuroinflammation is a key pathological mechanism in almost all acute and degenerative diseases in both the adult and in the developing brain(Hagberg et al 2012, Perry et al 2010, Prinz et al 2011). Microglia (MG) and infiltrating macrophages (Mφ) are major contributors to neuroinflammation(Perry et al 2010). These cells are capable of acquiring numerous complex functional states dependent on the specific nature of an insult or injury, including cytotoxic responses, immune regulation, or injury resolution(Wolf et al 2017). Limiting the cytotoxic activation of MG/Mφ while promoting injury resolution represents a rational neuroprotective strategy(Miron et al 2013, Rivest 2009), that requires an in-depth understanding of the molecular mechanisms controlling their phenotypes across forms of injury. Understanding of these regulatory mechanisms is increasing at an extraordinary rate for adult disorders. However, recent work has shown that MG do not reach maturity until around the time of birth in humans, or the equivalent developmental time point of approximately postnatal day 14 in rodents(Bennett et al 2016, Butovsky et al 2014, Krasemann et al 2017, Miyamoto et al 2016). These MG have specialised functions in brain development, such as synaptic pruning(Delpech et al 2016, Paolicelli et al 2011), but there is a striking paucity of knowledge on the regulators of MG phenotype in that is relevant to injury/insult to the developing brain.

Preterm birth is the commonest cause of death and disability in children under 5 years of age, exceeding deaths from malaria or pneumonia(Lim et al 2012). Preterm birth, birth before 37 of 40 weeks’ gestation, occurs in fifteen million infants yearly, rates are increasing in developed countries (e.g. 7% of all births in the UK; 13% in the US) and we are limited in our predictive and therapeutic options for these vulnerable infants. Up to 60% of infants born preterm we be left with persistent cognitive and neuropsychiatric deficits including autism-spectrum, attention-deficit disorders and epilepsy(Delobel-Ayoub et al 2009, Wood et al 2000). Insights into the previously poorly understood pathophysiology of preterm birth(Back et al 2005, Billiards et al 2008, Delobel-Ayoub et al 2009, Moore et al 2012, Verney et al 2012) have enabled us to design animal models of improved relevance to the constellation of injuries seen in contemporary cohort of infants, collectively called encephalopathy of the premature infant: MG/Mφ activation, oligodendrocyte maturation blockade, myelin deficits and axonapathy. MG/Mφ activation driving neuroinflammation in preterm born infants is often caused by exposure to maternal fetal infections or inflammation, such as chorioamnionitis, or post-natal sepsis. The endemic but often clinically silent maternal inflammatory events are considered the chief cause of preterm birth(Wu et al 2009) and exposure to inflammation is the strongest predictor of poor neurological outcome(Dammann & Leviton 2004, Hillier et al 1993, Wu et al 2009).

The Wnt (i.e. wingless-type MMTV [mouse mammary tumour virus] integration site) pathways have well studied critical roles in the early developmental events of body axis patterning, and brain cell fate specification, proliferation and migration(Sokol 2015). Wnt signalling is divided into the canonical Wnt/β-catenin and non-canonical Wnt/planar cell polarity (PCP) and Wnt/calcium pathways. Aberrant regulation of canonical WNT/β-catenin signalling is strongly implicated in the onset and progression of numerous cancers(Zhan et al 2017). The effects of Wnt pathway activation are highly cell and context specific and any key regulatory role of the Wnt pathways in the response of MG/Mφ to injury or insult has not been demonstrated, especially in the developing brain. Although, activation of the Frizzled receptors (a family of G protein-coupled receptors) with Wnt ligands has complex effects on some facets of MG activity *in vitro*(Halleskog et al 2012, Halleskog & Schulte 2013). Experimental evidence in other cell types also supports that the Wnt/ β-catenin pathway is involved in aspects of inflammatory signalling, namely there are links between expression of the inflammatory mediator COX2 (cyclooxygenase-2) and Wnt/β-catenin in cancer(Buchanan & DuBois 2006), and that β-catenin regulates at numerous levels the cytokine/pathogen induced activation of NF-κB (Ma & Hottiger 2016). As such, in the hunt for a regulator of immature MG activation state with a view to therapy design the Wnt pathway is an interesting and rationale candidate.

In the present study, we used our model of encephalopathy of prematurity and we have clearly demonstrated a causal link between MG/Mφ activation and oligodendrocyte injury/myelin deficit. Using a combination of approaches across species, including human, we reveal for the first time that inhibition of the Wnt/β-catenin pathway is necessary and sufficient to drive this MG/Mφ pro-inflammatory phenotypic transformation. We also verified the clinical relevance of the Wnt pathway to anatomical white matter structure in a cohort of human preterm born infant brain using an integrated imaging genomics analysis. Finally, we employed a novel 3DNA nanocarrier that carries cargo across the blood brain barrier (BBB) following non-invasive intraperitoneal (i.p.) administration and that then delivers cargo specifically to MG/Mφ. Using this 3DNA nanocarrier we show that preventing Wnt pathway down-regulation specifically in MG/Mφ reduces the pro-inflammatory MG/Mφ phenotype, white matter damage and long-term memory deficit in our model of encephalopathy of prematurity.

Altogether, these findings identify the Wnt/β-catenin pathway as a key regulator of MG/Mφ activation state across species and a potential target for the treatment of encephalopathy of prematurity and other neuroinflammatory conditions.

## Results

### MG activation drives hypomyelination in our model of encephalopathy of the premature infant

To understand the mechanisms underpinning encephalopathy of the premature infant in contemporaneous cohorts we previously set up a model mimicking its neuropathological, behavioural and imaging phenotypes(Favrais et al 2011, Krishnan et al 2017). Of note, the neuropathological similarities between our model and clinical studies includes MG/Mφ activation and hypomyelination due to the maturation blockade of oligodendrocytes but not cell death(Billiards et al 2008, Verney et al 2010, Verney et al 2012). Specifically, intraperitoneal (i.p.) injections of interleukin (IL)-1β (10µg/kg) were administered to mouse pups twice daily on postnatal days (P) 1-4 and once at P5 (**Figure 1a**). P1-P5 in the mouse is a developmental period roughly corresponding to 22-32 weeks of human pregnancy(Marret et al 1995) and this is the greatest period of vulnerability for white matter damage in preterm born infants. Exposure to systemic IL-1β recapitulates the systemic inflammatory insult of maternal/fetal infections, such as chorioamnionitis, that are responsible for setting up neuroinflammation via the activation of endothelial cell IL-1 receptors. In our model, we verified a direct causal link between MG/Mφ activation and neuropathology by selectively killing pro-inflammatory MG/Mφ and observing a reduction in myelin injury. Specifically, we used a intracerebral injection of gadolinium chloride (GdCl_3_), which kills pro-inflammatory MG/Mφ via competitive inhibition of calcium mobilization and damage to the plasma membrane, as previously validated *in vivo* (Du et al 2014, Fulci et al 2007, Miron et al 2013). This approach was chosen as ablation of MG with a Tamoxifen driven transgenic or other pharmacologic approaches (i.e. PLX3397, ganciclovir) is not feasible as our studies target maturing oligodendrocytes (as found in the preterm infant brain) that are present from P1. Also, embryonic depletion itself alters brain development(Squarzoni et al 2014). GdCl_3_ (200nmol) was injected into the corpus callosum of P1 mice prior to i.p injection of IL-1β or PBS. In the corpus callosum, 3 hours after the first IL-1β injection (at P1), immunohistological analysis revealed that the majority of MG/Mφ were IBA1+/COX2+ and the numbers of these cells was reduced by approximately 50% by GdCl_3_ (**Figure 1b,c**). This GdCl_3_-mediated reduction in MG/Mφ prevented the typical loss of myelin basic protein (MBP) in the corpus callosum at P30 (**Figure 1d,e**). A link between microglial activation and oligodendrocyte injury can also be recapitulated with IL-1β exposure of mixed glial cultures containing oligodendrocyte progenitor cells (OPCs), MG and astrocytes. In these *in vitro* conditions, there in reduced MBP+ cell density, without affecting the total number of oligodendrocytes (**Figure 1f)**. Exposure of pure OPC cultures to IL-1β in the absence of microglia or astrocytes did cause hypomyelination (data not shown). Altogether, these results validate in our model a causal link between MG/Mφ activation and the hypomyelination that is a hallmark of encephalopathy of prematurity.

**Figure 1.**
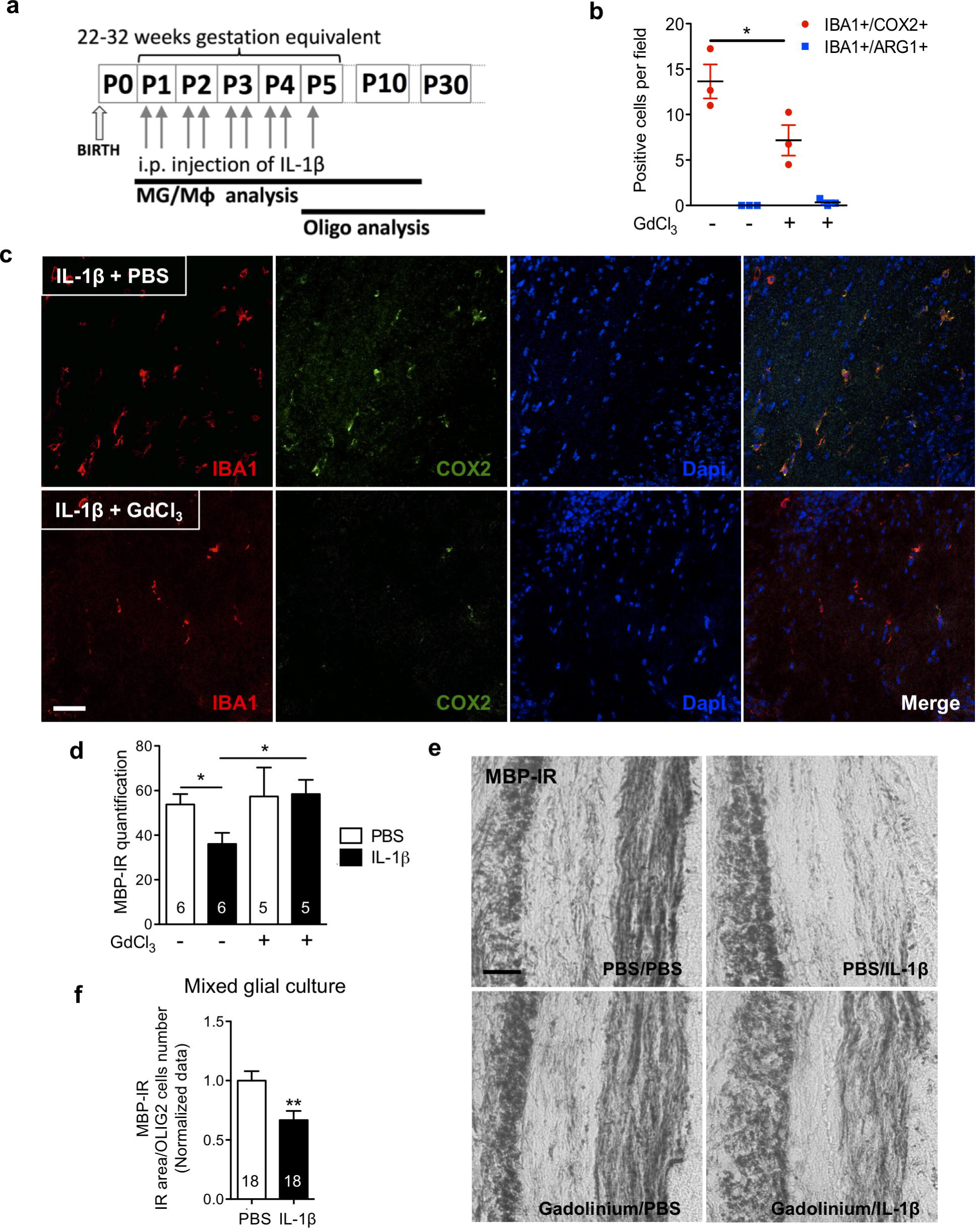
MG/Mφ are required to induce hypomyelination in our model of encephalopathy of prematurity. In (**a**) a schematic of the experimental paradigm for modelling encephalopathy of prematurity and also indicating the timing of analysis of MG/Mφ and oligodendrocytes (Oligo). Of note, experiments need to be performed between postnatal day (P1) and P5 as this is when oligodendrocyte maturation in the mouse matches that found in vulnerable preterm born infants, those born from 22-32 weeks’ gestation. In (**b-e**) we demonstrate that oligodendrocyte injury in our model is dependent on activated MG/Mφ by inducing cell death in these cells. Specifically, PBS or IL-1β exposed mice were injected in the corpus callosum with vehicle (PBS) or gadolinium chloride, GdCl3; 200 nmol. Quantification in (**b**) of the reduced number of pro-inflammatory (IBA1+/COX2+ cells) and stable numbers of anti-inflammatory (IBA1+/ARG1+ cells) MG/Mφ in corpus callosum at P3 (n=3/group, mean±SEM, Mann-Whitney test, * p<0.05). Representative images in (**c**) of immunoreactive staining in the corpus callosum for IBA1, COX2 and Dapi in mice at P3 treated as described for (a) and (b) (scale bar: 100µm). Following GdCl3 treatment, at P15 in (**d**) quantification of the reduced myelin injury as shown with MBP-IR by grey density in corpus callosum (mean±SEM, Mann–Whitney test, * p<0.05, n given inside bar on graph), and in (**e**) representative images showing MBP immuno-reactivity (MBP-IR) (scale bar: 20µm). The requirement for the presence of MG/Mφ to illicit oligodendrocyte demyelination was also verified *in vitro* in mixed glial cultures. Oligodendrocyte differentiation was shown to be decreased by IL-1β exposure by analysis of MBP-IR area normalized by OLIG2+ cell number in (**f**), where mean±SEM, ** p<0.01 and n are given inside each bar on graph.

### MG activation is persistent with specific temporal patterns in our model of encephalopathy of the preterm infant

We comprehensively characterized the morphology, gene and protein expression of MG/Mφ over time in our model to assess the relationship between activation states and injury. Our morphological analysis of GFP+ MG/Mφ from CX3CR1^GFP/+^ mice revealed a moderate but significant reduction of complexity (arborisation) at P3 when comparing MG/Mφ from IL-1β versus PBS injected animals but no difference in process length or density, cell body area, numbers of primary, secondary and tertiary processes, or the area covered by the MG processes in 2D (**Figure 2a,b**). Cell morphology was measured at 3 hours after the first inflammatory challenge (at P1) and at P3 when animals had been exposed to >48 hours of systemic inflammation driven neuroinflammation in vibratome cut sections using a custom designed script developed with the Acapella™ image analysis(Verdonk et al 2016).

**Figure 2.**
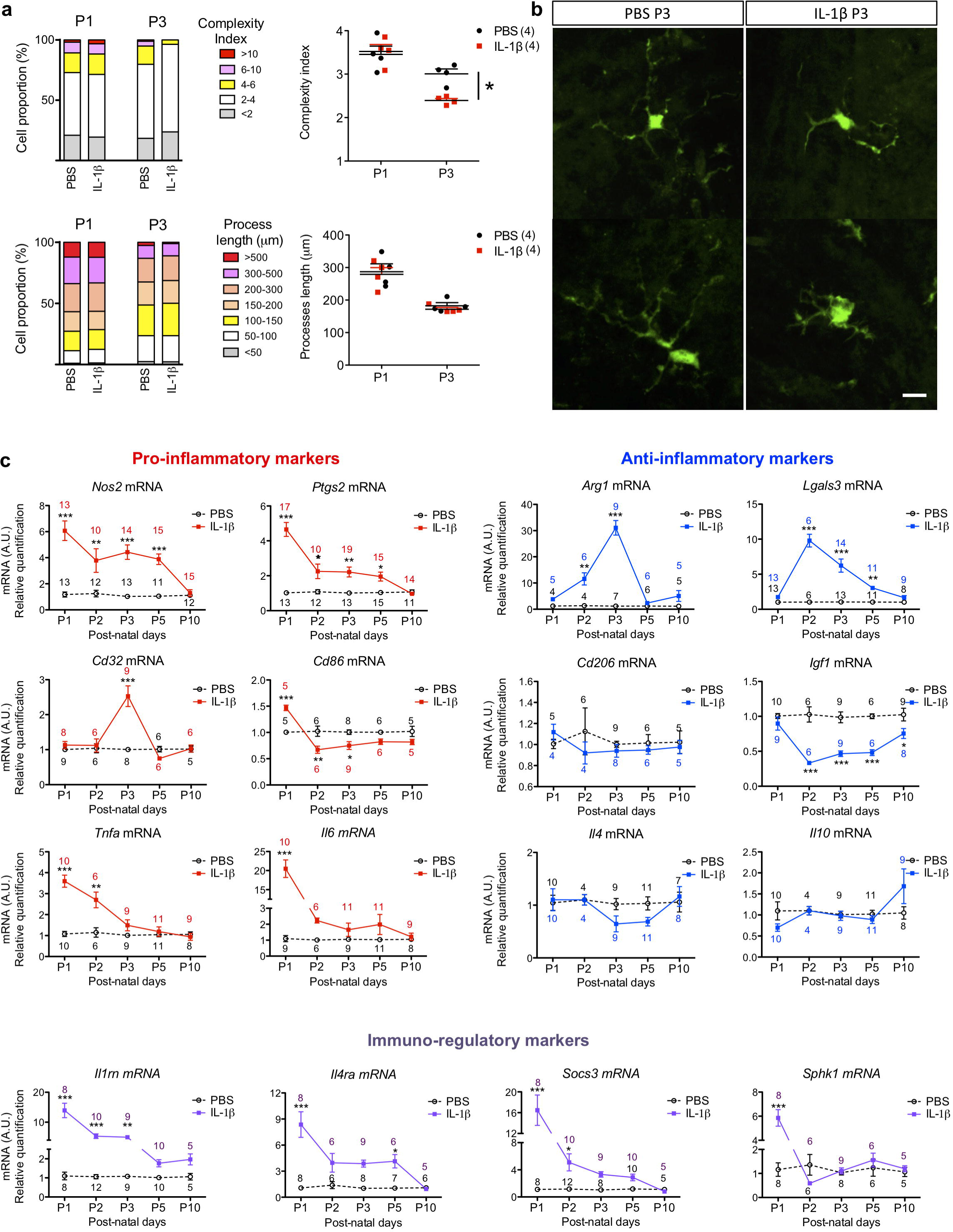
MG/Mφ have a subtly altered morphology and bi-phase alterations to phenotype in our model of encephalopathy of prematurity. Quantification in (**a**) of the complexity of MG/Mφ ramifications and process length in CX3CR1^GFP/+^ mice exposed via i.p. injection to PBS or IL-1β for 3 hours (P1) or for 48 hours (P3). The proportion (%) and absolute quantification of changes in MG/Mφ are shown; n=4/group, mean±SEM, Mann–Whitney test, * p<0.05. (**b**) Representative images of IL-1β induced decrease in the complexity index in GFP+ MG/Mφ at P3. Scale bar: 25µm. MG/Mφ phenotype over time via quantification in (**c**) of pro-inflammatory markers (*Nos2, Ptgs2, Cd32, Cd86, Tnf*α, and *Il6* mRNA), anti-inflammatory markers (*Arg1, Lgals3, Cd206, Igf1, Il4* and *Il10* mRNA) and immuno-regulatory markers (*Il1rn, Il4ra, Socs3* and *Sphk1* mRNA) by RT-qPCR in CD11B+ MG/Mφ magnetically sorted from brain in PBS or IL-1β injected mice. See also **Supp Figure 1**. mRNA expression as mean±SEM of the fold change relative to PBS group. Analysis made with a two-way ANOVA with *post hoc* Bonferroni‘s test (* p<0.05, ** p<0.01, *** p<0.001) and n for each data point is shown on the graphs in corresponding colours.

We further studied MG/Mφ activation using expression analysis of sixteen genes and six protein markers previously associated with polarized expression states(Chhor et al 2013, Miron et al 2013) at 5 time points. Gene expression analysis was performed in CD11B+ cells isolated by magnetically activated cell sorting (MACS, Miltenyi Biotec) from PBS- or IL-1β-exposed mice as per our previous studies(Chhor et al 2017, Krishnan et al 2017, Shiow et al 2017). Prior FACS analysis has demonstrated that the ratio of MG to other CD11B+ cells in this isolate is greater than 50 to 1(Krishnan et al 2017). Nevertheless, because we cannot exclude the presence of Mφ completely we use MG/Mφ as the descriptor for these cell isolates. Also, the purity of the MACS isolation was confirmed as >95% CD11B+ MG/Mφ by FACS and qRT-PCR, also as previously(Krishnan et al 2017, Schang et al 2014). qRT-PCR based gene expression analysis (**Figure 2c**) and immunofluorescence (**Supp Figure 1a-d**) showed that systemic exposure to IL-1β induced a robust pro-inflammatory MG/Mφ activation state at P1, only 3 hours following the first injection of IL-1β. Specifically, 5/6 markers of a pro-inflammatory state were significantly increased including *Il6* increased by 20-fold, and a 5-fold increases in *Ptgs2* and *Nos2*, all p<0.001. Of note, *Ptgs2* and *Nos2* remained elevated for 5 days, but *Il6* and *Tnfa* were returned to normal by P3. There was a similar robust increase in markers of immune-regulation at P1 as 4/4 markers were increased including an 8-fold increase in *Socs3* and >10-fold increase for *Sphk1* and *Il1rn*, all p<0.001. Expression of markers associated with an anti-inflammatory activation state were the most effected in animals exposed to systemic IL-1β exposure at P2 and P3, which is 48 or 72 hours after the start of IL-1β injections. These findings of a temporal change in MG/Mφ activation profiles were corroborated with immunohistological analysis of protein levels (**Supp Figure 1a-d**).

By combining our observations of morphology, gene and protein changes in MG/Mφ after systemic IL-1β exposure we have characterized a persistent state of activation responsible for injury to the developing white matter.

### The Wnt pathway is comprehensively down-regulated in activated MG in vivo and in vitro

In mouse pups exposed to systemic IL-1β we have characterized a persistent activation of MG between P1 and until at least P5, which induces a partial blockade of OPC maturation and subsequently hypomyelination(Favrais et al 2011). Using gene and predicted protein network-based analysis, we sought to identify molecular pathways regulating this MG/Mφ activation driving OPC injury. We hypothesized that the best time to pick-up the potential disrupting regulator pathways specific to developing MG was P5 when MG are activated and when the first effects of OPCs have been observed and P10 when MG activation is apparently resolved but injury to OPCs is well established. As such, we undertook genome-wide transcriptomics analysis of MG/Mφ (CD11B+) and OPCs (O4+) MACSed at P5 and P10 from the brains of mice exposed to IL-1β or PBS. The purity of the O4+ MACS isolation was verified with qPCR(Schang et al 2014) as outlined for CD11B fractions above. Genes showing differential expression between time points, P5 and P10, were identified for each cell-type, MG/Mφ (CD11B+) or OPCs (O4+), within each condition, IL-1β or PBS. Gene co-expression networks were inferred for each of the four sets of differentially expressed genes using Graphical Gaussian Models (GGMs)(Opgen-Rhein & Strimmer 2007). GGMs use partial correlations to infer significant co-expression relationships. We applied a False Discovery Rate (FDR) of <1% between any microarray probe pair in each of the four sets of differentially expressed genes sets previously identified: PBS-OPC, IL-1β-OPC, PBS-MG/Mφ and IL-1β-MG/Mφ. These four co-expression networks were found to exhibit unique topologies and functional annotations (**Supp Figure 2a,b; Supp Tables 1 and 2**). Compared to the three other conditions networks, the IL-1β exposed MG/Mφ co-expression network was the most highly interconnected and the densest indicating a strong co-regulation of gene expression under this condition (**Supp Figure 2a,b)**. Biological pathway analysis using the Broad Institute MsigDB database(Subramanian et al 2005) indicated enrichment for Wnt signalling pathway genes within the IL-1β exposed MG/Mφ network (FDR=8.5 ×; 10^−7^; **Figure 3a**; **Supp Table 2**). Due to the proposed importance of the Wnt pathway in OPC maturation(Fancy et al 2009) we undertook a specific analysis of data from the O4+ OPCs from mice subjected to systemic exposure to IL-1β. Analysis of microarray data from the O4+ OPCs did not show any significant enrichment for Wnt signalling genes in the gene co-expression response (**Supp Table 2**). This data highlight two important facts about Wnt dysregulation, firstly, that Wnt dysregulation is likely not a mechanism of oligodendrocyte maturation arrest in this model, and second, that changes in Wnt are not simply ubiquitous in the developing CNS in response to inflammation. Returning to the analysis of MG/Mφ, to highlight the directionality of the data, enrichments for Wnt signalling pathway sets were specifically noted among the genes significantly down-regulated by IL-1β exposure at P5 and P10 (**Supp Table 3**, see Methods). The known interactions among the Wnt pathway genes plus predicted interactors were then retrieved from an extensive set of functional association data by the GeneMania tool (Warde-Farley et al 2010) and we represented this sub-network (**Figure 3b**). We have also annotated this figure to highlight that the vast majority of targets predicted to interact in this model have down-regulated expression. Finally, qualitative gene profiling with qRT-PCR in CD11B+ MG/Mφ confirmed the down-regulation of mRNA expression for numerous Wnt pathway members including multiple Fzd receptors, *Lef1* and *Ctnnb1* in MG/Mφ from IL-1β-exposed mice (**Figure 3c).** The down-regulation of *Ctnnb1* mRNA was also accompanied by a significant decrease in the expression of β-catenin protein in CD11B+ MG/Mφ at P3 corresponding to the time point when the greatest morphological changes and pro-inflammatory gene activation is observed (**Figure 3d**).

**Figure 3.**
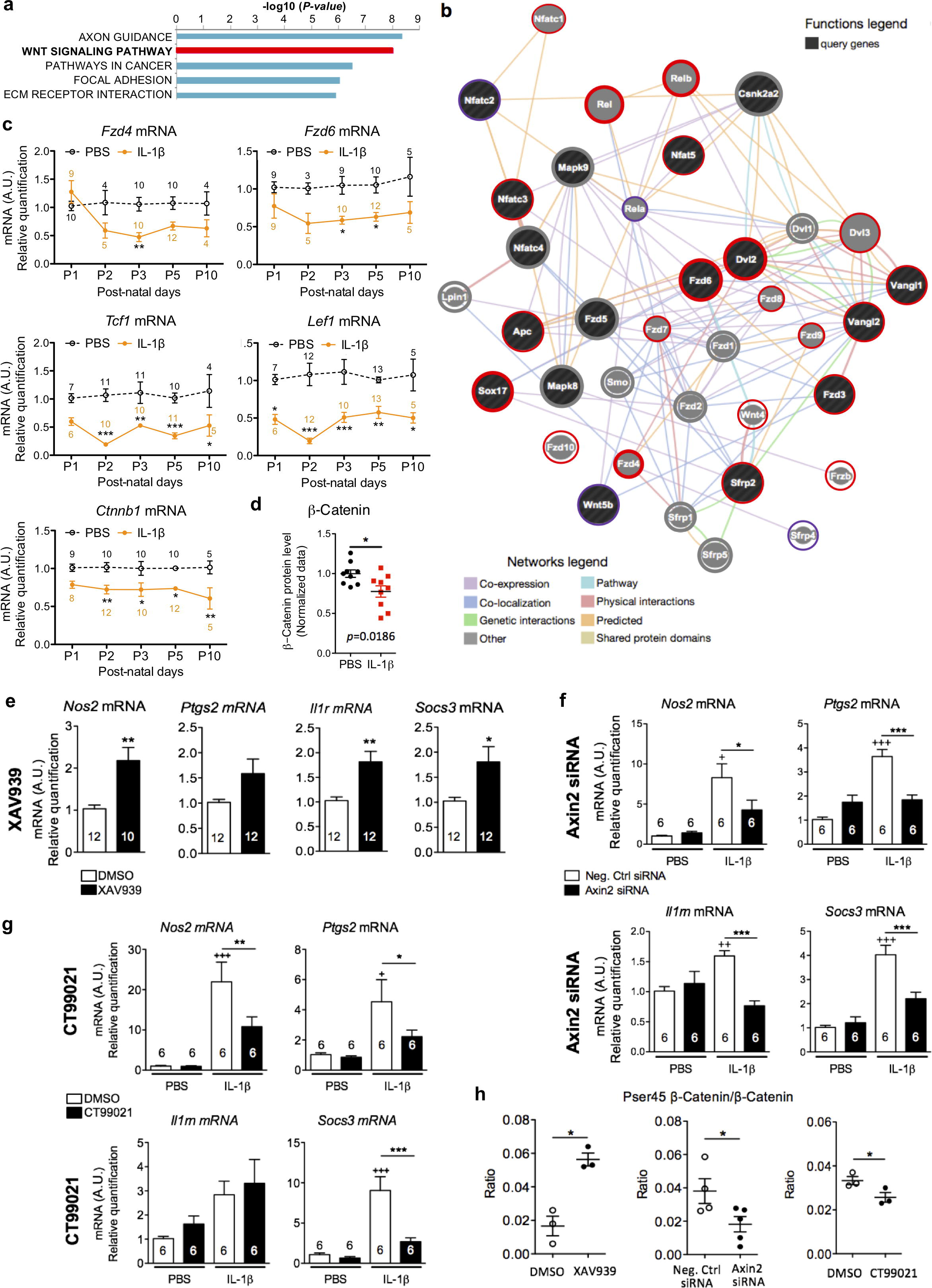
The Wnt pathway is down-regulated in MG/Mφ isolated from our model of encephalopathy of prematurity and down-regulation is necessary and sufficient for MG activation *in vitro*. Analysis of microarray data from MACS purified CD11B+ MG/Mφ from IL-1β and PBS administered mice was undertaken. This revealed (**a**) that changes in the Wnt signalling pathway are strongly significantly associated with MG/Mφ activation, see also **Supp Figure 2**. In (**b**) a cohesive network of multi-modal interactions between Wnt pathway genes in the MG/Mφ co-expression network can be generated via analysis using the GeneMania tool network (Warde-Farley et al 2010). Annotation of this network with a bold red circle indicates that the gene is down-regulated in the microarray data at P5 and P10, genes with a finer red line are down-regulated at a single time point and those in purple display a switch, being significantly higher at one and significantly lower at the other time point. In (**c**) validation of robust down-regulation in MG/Mφ of mRNA expression of *Ctnnb1, Tcf1, Lef1* and *Fzd 4,6* mRNA by RT-qPCR in CD11B+ MG/Mφ MACSed from brains of PBSor IL-1β-exposed mice at P1, 2, 3, 5 and 10. See also **Supp Figure 3**. Analysis made with a two-way ANOVA with *post hoc* Bonferroni‘s test: * p<0.05, ** p<0.01 *** p<0.001, numbers of samples for each data point shown on the graphs in corresponding colours. In (**d**) quantification of β-catenin by ELISA in CD11B+ MG/Mφ MACSed from brains of PBS or IL-1β-exposed mice at P3. β-catenin protein levels were reduced and are presented as a fold change relative to PBS-injected mice, (mean±SEM). Analysis made with a Student‘s t-test, * p<0.05. In (**e**) XAV939, a β-catenin pathway inhibitor induced a pro-inflammatory MG activation as assessed with quantification of *Nos2, Ptgs2, Il1rn* and *Socs3* mRNA by RT-qPCR in primary MG exposed to XAV939 (1µM) or DMSO for 4 hours. mRNA levels are presented as a fold change relative to DMSO group, mean±SEM, Student‘s t-test: * p<0.05, ** p<0.01. In (**f**) a reductions of pro-inflammatory MG activation by gene silencing using siRNA for Axin2 (30µM), a β-catenin inhibitor, or scrambled control (Ctrl) siRNA transfected into MG for 48 hours prior to 4 hours of IL-1β exposure, see also **Supp Figure 3h.** siRNA for Axin2 reduces IL-1β-induced *Nos2, Ptgs2, Il1rn* and *Socs3* mRNA. Data are presented as a fold change relative to negative siRNA group, mean±SEM, one-way ANOVA with *post hoc* Newman-Keuls‘s test: effects of IL-1β on gene expression are shown with + p<0.05, ++ p<0.01, +++p<0.001; effects of Axin2 siRNA to alter the IL-1β effects are shown with * p<0.05, ** p<0.01, ***p<0.001. In (**g**) Inhibition of GSK3β, aβ-catenin inhibitor, by CT99021 (1µM) treatment 1 hour prior to 4 hours of IL-1β exposure reduces IL-1β-induced *Nos2, Ptgs2, Il1rn* and *Socs3* mRNA overexpression. mRNA levels are quantified by qRT-PCR and presented as a fold change relative to control/DMSO group, (mean±SEM, One-way ANOVA with *post hoc* Newman-Keuls‘s test: + p<0.05, ++ p<0.01, +++p<0.001 indicate a statistically significant difference between PBS+DMSO and IL-1β+DMSO; * p<0.05, ** p<0.01, ***p<0.001. In (**h**)effects of modulating Wnt with XAV939, Axin2 siRNA and CT99021 on phosphorylation of β-catenin(PSer45 β-catenin/β-catenin ratio) were quantified by ELISA in MG. XAV939 (1µM) inhibits β-catenin while CT99021 (1µM) or siRNA against *Axin2* (30µM) activate it when you compare to their respective vehicle (DMSO or negative control siRNA). Data are expressed as the mean±SEM (*p<0.05 in a Mann-Whitney test). For all bar graphs the n for each data point is indicated on or above the bar.

To extend this *ex vivo* CD11B+ cell data and set up an *in vitro* model for further study, qRT-PCR and immunofluorescence analysis of mouse primary cultured MG was undertaken. The cultures are >97% pure for MG as validated with IHC and qRT-PCR, as previously(Chhor et al 2013). Exposure of MG to IL-1β *in vitro* induces a similar pro-inflammatory phenotype as is found in MG/Mφ from mice exposed to systemic IL-1β (**Supp Figure 3a,b**). Furthermore, we observed a similar down-regulation of Wnt pathway members including mRNA encoding *Fzd* receptors, *Ctnnb1* and *Tcf1* (**Supp Figure 3c**). Inducing a pro-inflammatory activation in MG *in vitro* also induced an upregulation of mRNA encoding *Axin1* and *Axin2*, two specific inhibitors of β-catenin nuclear translocation (**Supp Figure 3c**) and reduced the protein levels for β-catenin as assessed with immunofluorescence (**Supp Figure 3d**), and ELISA (**Supp Figure 3e**). There was no significant change in the ratio of phosphorylated (Pser45) β-catenin to β-catenin (**Supp Figure 3e**). In contrast, IL-4, a classic anti-inflammatory stimulus, increased Wnt/β-catenin activation as indicated by strongly increased *Lef1* mRNA and decreased the Pser45 β-catenin/β-catenin ratio measured via ELISA (**Supp Figure 3f,g**).

Altogether these data demonstrate that a pro-inflammatory MG/Mφ phenotype is associated with a robust and comprehensive down-regulation of the gene and protein expression of members of the Wnt/β-catenin signalling pathway *in vivo* and *in vitro*.

### Wnt/β-catenin pathway activity negatively correlates with pro-inflammatory MG activation in vitro

We next sought to determine if Wnt/β-catenin pathway modulation alone is sufficient to drive phenotypic changes in MG *in vitro*. Pharmacological inhibition of the Wnt/β-catenin pathway *in vitro* with XAV939, a tankyrase inhibitor that stabilizes the Wnt/β-catenin pathway inhibitor Axin2 (Fancy et al 2011), was sufficient to promote a pro-inflammatory phenotype, mimicking the effects of IL-1β (**Figure 3e**). Conversely, the IL-1β-induced pro-inflammatory MG phenotype was blocked by activating the Wnt pathway using siRNAs against Axin2 (**Figure 3f;** validation of siRNA efficacy **Supp Fig 3h**) or a pharmacological inhibitor of GSK3β that phosphorylates and inactivates β-catenin, CT99021(Ring et al 2003) (**Figure 3g**). As expected, we verified with ELISA that the Pser45 β-catenin/β-catenin ratio was increased with XAV939, and decreased with *Axin2* siRNA and CT99021 (**Figure 3h**). In contrast, we tested in PBS and IL-1β-exposed primary MG a non-isoform specific block of PKC signalling with Chelerythrine. Chelerythrine targets the non-canonical Wnt pathways and exposure of doses from 1-3μM had no effect on MG activation (**Supp Figure 4**).

**Figure 4.**
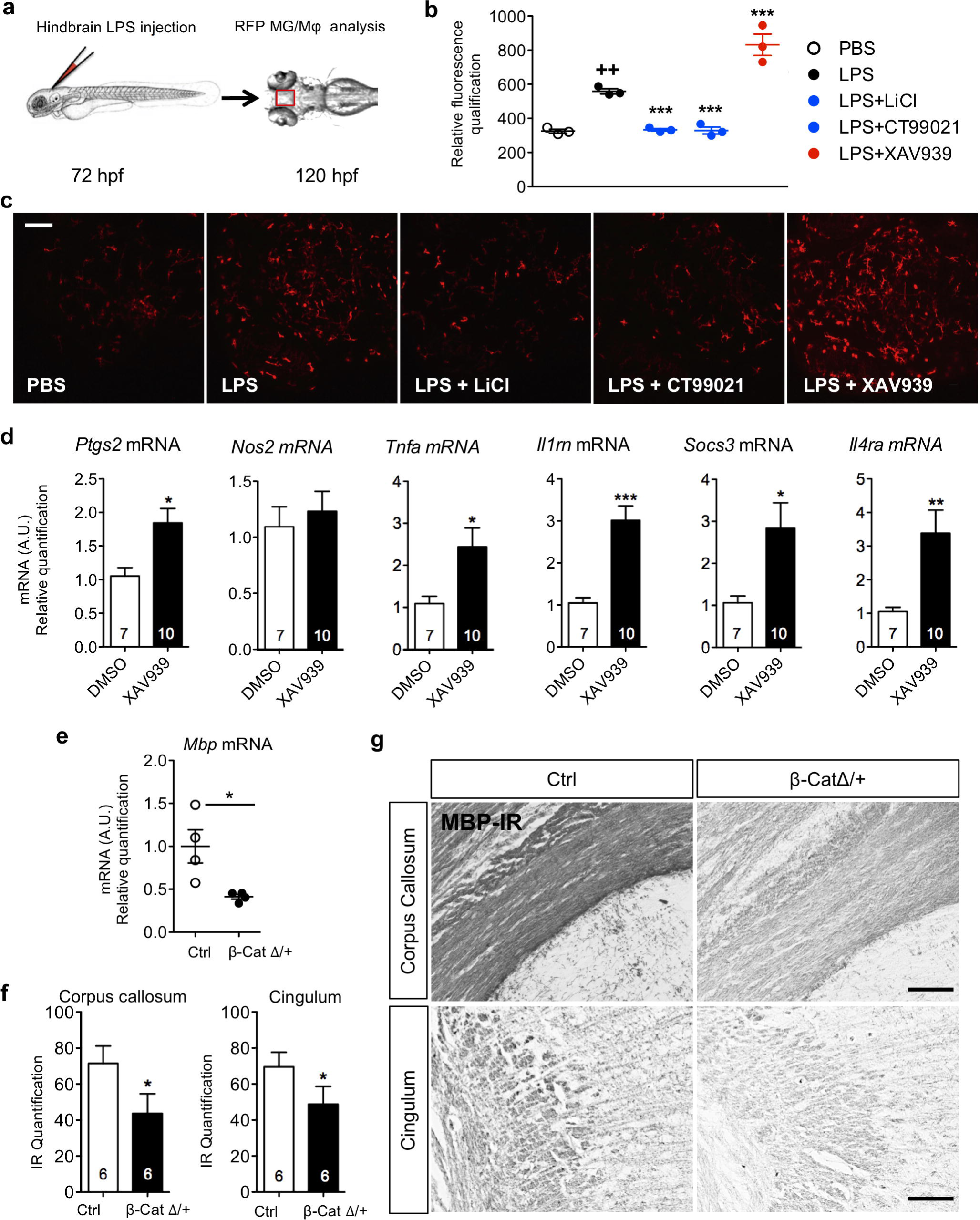
Wnt/β-catenin pathway regulates MG/Mφ activation and hypomyelination *in vivo*. Wnt signalling was modulated *in vivo* with pharmacological techniques in zebrafish (**a-c**) and mice (**d**) and using gene knockout in mice (**e-g**). In (**a**) LPS microinjection (5ng/injection) into the zebrafish hindbrain at 72 hpf in pu1::Gal4-UAS::TagRFP morpholinos followed by analysis at 120 hpf. In (**b**) quantification of RFP labelled MG and in (**c**) representative images of RFP MG in these zebrafish brains after LPS injection in the presence or not in their growth solution of modulators of Wnt/β-catenin signalling, LiCl (80mM), XAV939 (5μM), CT99021 (3μM). Agonists of Wnt (LiCl, CT99021) prevented LPS driven MG immunoreactivity and further antagonizing Wnt (XAV939) enhanced this inflammatory reactivity. Data are mean±SEM, One-way ANOVA with *post hoc* Newman-Keuls‘s test:** p<0.01, ***p<0.001. Scale bar: 40μm. In (**d**) analysis of CD11B+ cells isolated from P1 mice injected i.c.v. with XAV939 (0.5 nmol) alone demonstrating increased MG/Mφ activation. Quantification of *Nos2, Ptgs2, Tnf*α, *Il1rn, Socs3* and *Il4ra* mRNA by RT-qPCR. mRNA levels are presented as a fold change relative to PBS/DMSO i.c.v. injected mice. Data expressed as the mean±SEM, Student‘s t-test: * p<0.05, ** p<0.01 and ***p<0.001. In **(e-g)** β-catenin deficit in MG driven by β-catenin^flox/+^/LysM^Cre/+^ (β-cateninΔ/+) mice induces hypomyelination. In (**e**) quantification of *Mbp* mRNA by RT-qPCR at P10 in the anterior brain of β-catenin^Δ/+^ mice. mRNA levels arepresented as a fold change relative to WT mice, mean±SEM, Mann–Whitney test: * p<0.05. In (**f**) theβ-catenin deficit in β-catenin^Δ/+^ mice itself induces a decrease of myelin-binding protein MBP immunoreactivity (MBP-IR) in the corpus callosum and cingulum at P30 relative to Ctrl (β-catenin^Δ/+^/C57Bla6) mice. Results are expressed by grey density, mean±SEM, Mann–Whitney test: *p<0.05. In (**g**) representative images of MBP-IR in the corpus callosum and cingulum at P30 in Ctrl and β-catenin^Δ/+^. Scale bar: 60μm. For all bar graphs the n for each data point is indicated on or above the bar.

Altogether, these data verify in pure cultures of primary MG that that modulating the Wnt/β-catenin pathway is sufficient and necessary to effect MG activation state.

### Wnt/β-catenin pathway activity negatively correlates with pro-inflammatory MG activation in vivo

We further tested our working hypothesis that Wnt/β-catenin signalling drives a pro-inflammatory phenotype in MG/Mφ *in vivo* using non-cell specific approaches in zebrafish and mice and then transgenic mice with MG/Mφ specific β-catenin knock-down. In zebrafish larvae at 72 hours’ post-fertilization (hpf) that express RFP in MG/Mφ (Tg[pU1::Gal4-UAS-TagRFP]) we injected into the hindbrain the prototypical pro-inflammatory agent lipopolysaccharide (LPS) (**Figure 4a**). LPS was used due to the unavailability of recombinant zebrafish IL-1β and prior work demonstrating the expression by zebrafish of receptors for LPS, the Toll-like 4 receptor (TLR4), and the requisite downstream signalling cascade(van der Sar et al 2006). LPS-induced neuroinflammation was confirmed after 48 hours by a significant increase in fluorescent MG/Mφ in the optic tectum (**Figure 4b,c**). We observe a similar increase in MG/Mφ immunofluorescence in our mouse model with IBA1 and MAC1 immunofluorescence(Favrais et al 2011). As expected, exposure of zebrafish larvae to the Wnt/β-catenin antagonist XAV939 exacerbated the LPS-induced MG/Mφ activation, while conversely exposure to Wnt agonists acting via the canonical GSK3β dependent pathway with CT99021 or LiCl prevented the LPS induced MG/Mφ activation (**Figure 4b,c**).

Subsequently, we studied the specific relationship between Wnt/β-catenin pathway activity and MG/Mφ activation in mice using pharmacological and genetic approaches. Firstly, we inhibited the Wnt/β-catenin pathway by intracerebroventricular (i.c.v.) injection of XAV939 in P1 pups and isolated CD11B+ MG/Mφ 3 hours later. In agreement with our hypothesis, *in vivo* Wnt inhibition with XAV939 mimicked the effects of systemic IL-1β on gene expression in MG/Mφ (**Figure 4d**). Furthermore, to specifically target the Wnt pathway in MG/Mφ, we bred LysM^Cre/Cre^ with β-catenin^Flox/Flox^ mice to generate β-CatΔ/+ transgenic mice. Recombination, and as such KO, of β-catenin occurs in ≥20% of MG/Mφ in this Cre line(Cho et al 2008, Degos et al 2013, Derecki et al 2012). In the absence of external inflammatory stimuli, β-CatΔ/+ mice with MG expressing reduced levels of β-catenin developed a myelin deficiency that mimicked the damaging effects of systemic IL-1β exposure in our encephalopathy of prematurity model. Specifically, these mice had reduced *Mbp* mRNA in the anterior cortex at P10 (**Figure 4e**) and reduced MBP immunoreactivity in the corpus callosum and cingulum at P30 (**Figure 4f,g**).

Collectively, these data show that in an invertebrate and a vertebrate species that down-regulation of the Wnt pathway is sufficient and necessary to drive a pro-inflammatory MG/Mφ activation state. Of note, even a partial MG/Mφ-specific β-catenin deletion *in vivo* in the absence of any external stimuli is sufficient to induce MG/Mφ activation leading to a myelination defect.

### The Wnt pathway regulates activation state in primary human MG in vitro

We then verified a similar relationship between decreased Wnt and increased pro-inflammatory activation in primary human MACSed CD11B+ MG/Mφ isolated from cerebral tissues collected from scheduled terminations at 19 and 20 weeks’ gestational age. Based on the scarcity and value of these cells we chose to use LPS to induce a pro-inflammatory MG response based on previous studies(Melief et al 2012). Human primary MG/Mφ were activated to a pro-inflammatory state as indicated by modified morphology (**Figure 5a**) and increased COX2 gene expression (*PTGS2* mRNA) (**Figure 5b**). In agreement with what we observed in our mouse and zebrafish experiments these pro-inflammatory activated human primary MG had a reduction in their expression of mRNA for the β-catenin gene, *CTNNB1* (**Figure 5c**).

**Figure 5.**
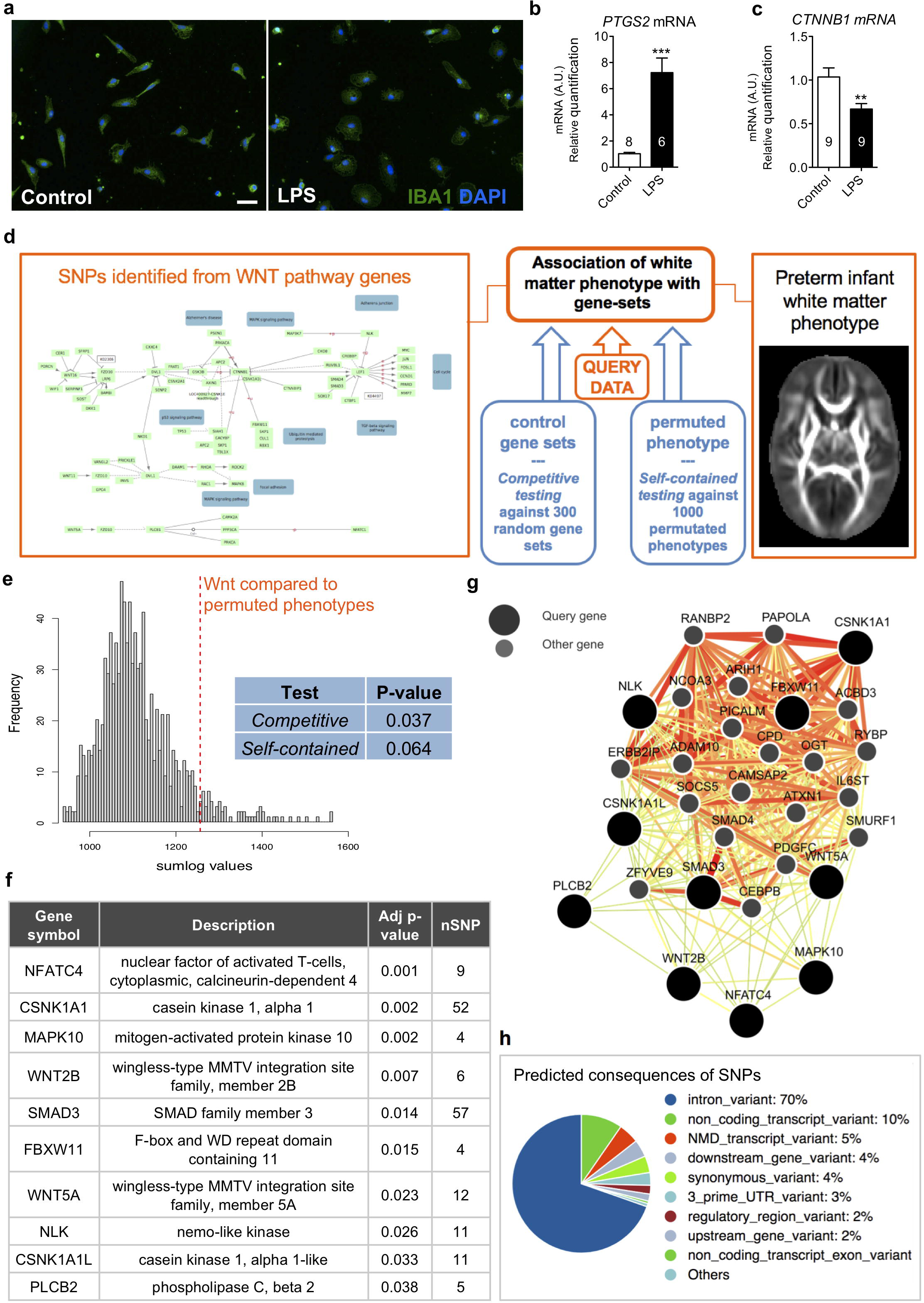
In human MG activation leads to Wnt/ β-catenin down-regulation and genetic variation in the Wnt pathway associates with preterm infant white matter phenotype. In (**a**) human primary MG exposed to PBS or LPS showing immunoreactive changes with IBA1 plus DAPI, scale bar = 50µm, and (**b**) mRNA expression for COX2 (*PTGS2)* verifying MG activation by LPS and (**c**) the associated decrease in the expression of the gene for β-catenin (*CTNNB1)*, t-test,**p<0.01, ***p<0.0001. In (**d**) a schematic of the analysis for any association between SNPs in the WNT pathway and the preterm infant white matter phenotype. Common genetic variation in the WNT gene-set was enriched for variants associated with tractography features when compared to random matched gene-sets, competitive test, p=0.037, 1000 permutations, (**e blue inset**) and is also associated with white matter probabilistic tractography in preterm infants, compared with the null background, self-contained test, p=0.064, 1000 permutations (**e blue inset, e**). In (**f**) are the functional descriptions of the ten genes within the WNT gene-set containing SNPs that were most significantly associated with the tractography phenotype. In (**g**) the relationships between these ten genes significantly associated with the tractography phenotype is reconstructed in a high confidence interaction network specific to the human brain, retrieved from known tissue-specific expression and regulatory data (Greene et al 2015). In (**h**) is the predicted consequences of all of the SNPs found in the top 10 ranked WNT genes shown in (f), a total of 42 SNPs.

### Genomic variance in WNT pathway genes is relevant to human preterm infant white matter structure

Our experimental data show that genetic knockdown of Wnt/β-catenin signalling is sufficient itself to drive brain injury in transgenic mice. In a conceptually similar manner, but with an expected much smaller effect size, we reasoned that that genetic variation in *WNT* pathway genes would be associated with surrogates of white matter mediated connectivity in preterm born infants. We hypothesize that, at least in part, any link between WNT variation and connectivity would be mediated by differences in the innate response to inflammatory challenge in these preterm infants. Although our genetic analysis is agnostic with respect to cellular mechanism or type, a link between WNT variations and brain structure would support further research into WNTs role in injury, and using WNT variants for preterm born infant injury/outcome stratification. To test our prediction of a link between WNT and preterm brain connectivity we performed an analysis where we jointly analysed imaging data and genomics data from 290 preterm born infants. This approach has previously uncovered novel genetic variants associated with brain connectivity phenotype (Krishnan et al 2017, Krishnan et al 2016). Specifically, we assessed single nucleotide polymorphisms (SNPs), DNA sequence variations where a single nucleotide varies between individual members of a species that can help pinpoint contributions of specific genes to disease states.

To define the intra-cerebral anatomical connectivity phenotype in our preterm cohort, we used 3-T dMRI and optimized probabilistic tractography as previously described(Robinson et al 2008, Shi et al 2011), adjusting the results for post-menstrual age at scan and at birth. Regions of interest for seeding tractography were obtained by segmentation of the brain based on a 90-node anatomical neonatal atlas(Shi et al 2011), focusing on cortico-cortical connections. DNA extracted from saliva was genotyped and a gene-set-of-interest defined using SNPs in the *WNT* signalling pathway (entry hsa04310) of the Kyoto Encyclopaedia of Genes and Genomes (KEGG) (*n*=141 genes).

SNPs were mapped to all of the genes in the KEGG WNT pathway using the JAG tool and Genome Reference Consortium Human Build 37 (GRCh37; hg19), using 2kb up/downstream, in line with NCBI practices (NCBI 2005). Following SNP-gene mapping, we tested the null hypothesis that variation in WNT pathway genes is not associated with the preterm tractography phenotype using Joint Association of Genetic Variants (JAG)(Lips et al 2015). Specifically, genes in the WNT-set were tested in two ways (**Figure 5d**). Firstly, we performed a “competitive/enrichment test” to determine whether the *WNT* pathway genes were no more associated with the connectivity phenotype than 300 randomly drawn non-*WNT*-pathway gene-sets. Secondly, we performed a “self-contained/association test” to determine whether there is a comparable association between the original connectivity phenotype and *WNT*, versus 1000 random permutations of the phenotype and *WNT*. The “competitive test”, in which we compared to randomly generated sets of genes, showed a significant enrichment of genetic variants associated with the phenotype in the *WNT* gene-set compared to random gene sets (empirical p=0.037, **Figure 5e inset**). The conservative “self-contained” test in which we compared to 1,000 permutated phenotypes indicated that the WNT signalling gene-set was associated with our phenotype; i.e., white matter probabilistic tractography features (empirical *p*=0.064; **Figure 5e histogram** & **inset**). We performed a gene-by-gene analysis of SNPs within the *WNT* signalling gene-set to investigate which genes might be predominantly contributing to the collective pathway enrichment signal. This test highlighted a group of 10 genes with individual significant association with the tractography phenotype (empirical *p*≤0.05, 1,000 permutations). These ten genes included 8 associated with Wnt/β-catenin signalling *NFATC4, CSNK1A1, MAPK10, WNT2B, SMAD3, FBXW11, NLK, CSNK1A1L*, and 2 associated with the Wnt/Ca^2+^ or Wnt/PCP pathways, *PLCB2* and *WNT5A* (**Figure 5f; Supp Table 4**). Although, we cannot determine specifically how these SNP variants would alter the brain phenotype we found using the Genome-scale Integrated Analysis of gene Networks in Tissues (‘GIANT’) tool that these genes created a coherent interaction network within a human brain-specific gene interaction network (**Figure 5g; Supp Table 5**). GIANT hosts searchable genome-scale functional maps of human tissues, integrating an extensive collection of experimental data sets from publications including expression, regulatory and protein data(Greene et al 2015). We also performed an exploratory investigation of function for the 42 SNPs from the 10 genes predominantly contributing to the collective pathway enrichment signal to provide information related to function. We used the Gene Effect Predictor within the Consortium Human Build 37 (GRCh37; hg19) tool to study the 42 SNPs, whose full details are given in **Supp Table 6**. All 42 SNPs were existing in the database and they were predicted to have 463 consequences on the genome (**Supp Table 7**). Specifically, these SNPs overlapped with 103 separate gene transcripts, 7 of them overlapped regulatory sequences and overall 95% were synonymous variants and 5% were missense variants (**Figure 5h).**

These linked imaging-genomic analyses demonstrate a relationship between genetic variation in the WNT pathway with an important clinical phenotype. They provide adjunct support for a causal relationship between the Wnt pathway and the brain phenotype that has been established with our experimental data.

### A Wnt pathway agonist delivered specifically to MG by a nanocarrier in vivo prevents neuroinflammation-mediated hypomyelination and the associated cognitive impairment

There is a paucity of neurotherapeutic options for damage to the preterm born infant brain and many other neuroinflammatory-mediated neurological disorders. We have clearly demonstrated a link between decreased Wnt/β-catenin pathway activation and a pro-inflammatory MG activation state. In addition, we have shown the relevance of this pathway in human MG and preterm infant brain structure. As such, we complete this study by demonstrating that the Wnt pathway is a viable therapeutic target. To achieve this, we specifically increased Wnt pathway activity in MG/Mφ using a peptide inhibitor of GSK3β, which has the effect of preventing the formation of the β-catenin degradation complex. We delivered this peptide to MG/Mφ *in vitro* and *in vivo* by conjugating it to a 3DNA nanocarrier together with a fluorescent tag (Cy3)(Genisphere, PA, USA). 3DNA nanocarriers are interconnected monomeric subunits of DNA of approximately 200nM in diameter that are intrinsically capable of endosomal escape(Muro 2014). In brief, we validated that our 3DNA-peptide conjugate had MG specificity and immunomodulatory efficacy *in vitro*, that is crosses the BBB, that it has cell specificity *in vivo,* and finally that it is neurotherapeutic *in vivo* in our systemic-inflammation (IL-1β) driven model of encephalopathy of the preterm born infant.

First, we demonstrated *in vitro* that 3DNA nanocarrier Cy3 tagged conjugates are specifically internalized by MG in mixed glial cultures (**Supp Figure 5a)** and by MG in primary cultures after 4 hours of incubation (**Supp Figure 5b**). When 3DNA was conjugated to both Cy3 and the peptide L803mts, an agonist of canonical Wnt signalling via inhibition of GSK3β, it was also specifically internalized by MG in primary culture (**Supp Figure 6a**). We also used the 3DNA nanocarrier tagged with Cy3 to deliver a cargo of L803mts to MG/Mφ that had been activated with IL-1β *in vitro*. Treatment of primary MG with L803mts delivered with 3DNA reversed the IL-1β-induced activation of the MG but had no effect on PBS-treated MG as assessed with gene expression analysis (**Supp Figure 6b**).

**Figure 6.**
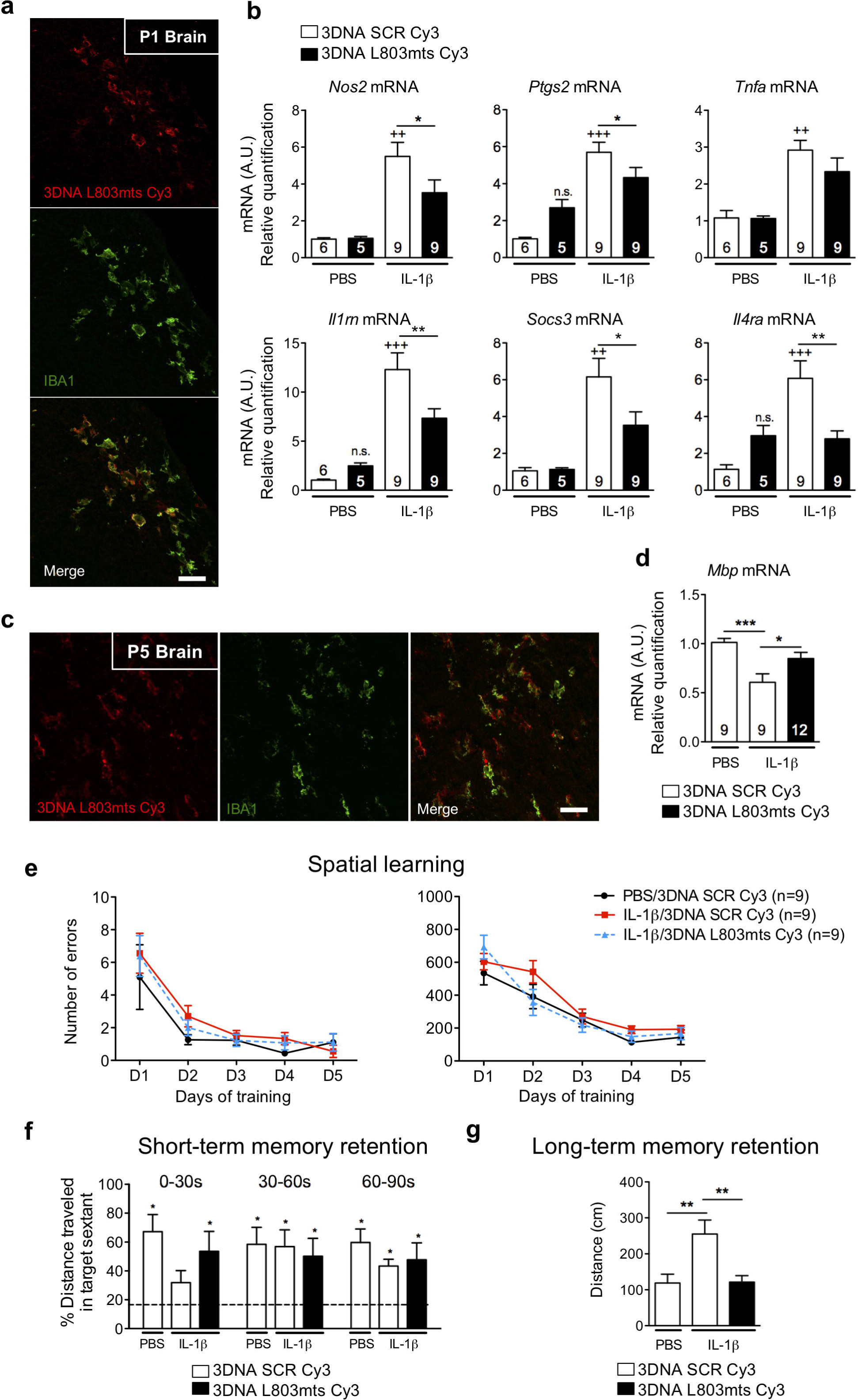
Nanocarrier delivery of a Wnt/ β-catenin pathway activator L803mts regulates MG/Mφ activation, and prevents hypomyelination and memory impairment in our encephalopathy of prematurity model. In (**a**) representative images of the *in vivo* uptake of 3DNA by IBA1+ cells in the subventricular white matter 4 hours after i.c.v. administration of 3DNA SCR Cy3 (vehicle) or 3DNA L803mts Cy3 (200ng), scale bar: 40μm. See also **Supp Figure 4a,b** and **Supp Figure 5**. In (**b**) effects of i.c.v. injection of 3DNA L803mts Cy3 to reduce the IL-1β-induced MG/Mφ activation as evaluated by qRT-PCR quantification of *Nos2, Ptgs2, Tnf*α, *Il1rn, Socs3* and *Il4ra* mRNA in CD11B+ MG/Mφ cells. mRNA levels are presented as a fold change relative to 3DNA SCR Cy3/PBS group (n for each data point is indicated on the bar, mean±SEM, One-way ANOVA with *post hoc* Newman-Keuls‘s test: ++ p<0.01, +++p<0.001 indicate a statistically significant difference between the PBS and the IL-1β groups also exposed to 3DNA SCR Cy3. For the effects of Wnt modulator delivery, * p<0.05, ** p<0.01. In (**c**) representative images of *in vivo* uptake of 3DNA by IBA1+ cells in subventricular white matter at P5 after i.p. injections of 3DNA L803mts Cy3 or 3DNA SCR Cy3 (500ng/injection) concurrently with PBS or IL-1β i.p. injections from P1-P5. Scale bar: 40μm. In (**d**) effect of i.p. injections of 3DNA L803mts Cy3 to prevent the IL-1β-induced loss of *Mbp* mRNA levels as quantified by qRT-PCR in anterior brain of mice at P10. mRNA levels are presented as a fold change relative to 3DNA SCR Cy3/PBS group (n for each data point is indicated on the bar, mean± SEM, one way-ANOVA with Bonferroni‘s post hoc test: ***p<0.001, * p<0.05). See also **Supp Figure 4c,d.** To show that systemically delivered 3DNA L803mts Cy3 led to functional improvements in our model together with the structural changes we performed Barnes maze testing to evaluate spatial learning (**e**) and memory (**f,g**) in 3-month-old mice after i.p. injections of PBS or IL-1β from P1-P5 concurrently with treatment with 3DNA SCR Cy3 or 3DNA L803mts Cy3. See also **Supp Figure 6**. In (**e**) learning was evaluated by recording the number of errors and distance travelled to reach the target (escape box) over 5 consecutive days. In (**f**) we show that 3DNA SCR Cy3 treatment ameliorated the negative effects of IL-1β-exposure on memory retention present in the 0-30 second trial period on day 5 of the test (probe trial) as measured by recording of the distance travelled in the target sextant, and in (**g**) we show that 3DNA SCR Cy3 treatment ameliorated the negative effects of IL-1β-exposure on the long-term memory deficits as measured via the distance travelled to reach the target on the 15^th^ day after the start of testing. For probe trial comparisons, a univariate t test was performed to compare the % time spent in the target sextant to the theoretical value 16.67%, i.e. when the mouse spent equal time within each sextant, * p<0.05. For long-term memory analysis, a one-way ANOVA with Bonferroni *post hoc was used \*\** p<0.01.

Moving *in vivo*, we aimed to generate the proof-of-concept that the 3DNA nanocarrier delivering L803mts and tagged with Cy3 can be taken up by MG/Mφ and influence MG/Mφ phenotype. Specifically, the L803mts and Cy3 conjugated 3DNA nanocarrier or the scrambled (SCR) peptide control and Cy3 conjugated 3DNA nanocarrier was injected i.c.v. into P1 mice, 1 hour prior to our standard i.p. IL-1β injection. Four hours after i.c.v. injection, we observed Cy3 fluorescence exclusively co-localized with IBA-1+ staining in the periventricular white matter (**Figure 6a**). In the same paradigm, using qRT-PCR analysis of isolated CD11B+ MG/Mφ we demonstrated that i.c.v. delivered 3DNA loaded with L803mts and tagged with Cy3 reversed the effect of IL-1β on the MG/Mφ phenotype but that the 3DNA-L803mts-Cy3 had no significant effect on MG/Mφ from PBS-treated mice (**Figure 6b**). As i.c.v. injections cause aggravated tissue injury we subsequently undertook a translationally orientated trial using i.p. delivery of Cy3-tagged 3DNA conjugated to L803mts. The 3DNA L803mts Cy3 or 3DNA SCR Cy3 control was injected i.p. daily between P1 and P5, in parallel with PBS or IL-1β. At P5, immunofluorescence in the anterior periventricular white matter confirmed that 3DNA L803 mts Cy3 was specifically taken up by IBA1+ MG/Mφ (**Figure 6c**). We also assessed uptake of our 3DNA conjugate by populations of macrophage in the body after i.p. delivery at P5. Using the same IHC techniques that clearly shows uptake in MG/Mφ we did not find Cy3 positive staining in the spleen or liver (**Supp Figure 5c,d**).

To validate the *in vivo* neurotherapeutic efficacy of our novel nanocarrier-mediated WNT agonist therapy, we assessed the IL-1β-induced myelination deficit by measuring *Mbp* mRNA. Treatment with i.p. 3DNA L803mts Cy3 significantly prevented the IL-1β-induced reduction in *Mbp* mRNA (**Figure 6d**). In order to evaluate whether this improvement in neuropathology with nanocarrier mediated treatment also reverses the behavioural deficits observed in adults in this paradigm, spatial learning and memory were assessed starting at P50. Basic behavioural parameters we first tested via spontaneous locomotion and rearing using actimetry and exploratory behaviour with an open field test. As expected no changes in these basic parameters was observed between groups of mice., i.e. controls (PBS+3DNA SCR Cy3), injured (IL-1β+SRC Cy3) or the treatment group (IL-1β+3DNA L803mts Cy3) (**Supp Figure 7a,b**). Spatial learning and memory was tested via the Barnes maze test, which assesses spatial learning over 5 days of probe trials plus short term memory retention at the end of the learning trials and long term memory retention 10 days later. Spatial learning (i.e., distance travelled and number of errors) itself was not modified by IL-1β exposure (**Figure 6e**). However, mice exposed to IL-1β had a short-term memory deficit based on a decreased distance travelled in the target sextant during the first 30 seconds of the probe trial on the final day of testing (**Figure 6f**). In addition, IL-1β exposure caused deficits in long term memory recall, as assessed as the increased distance travelled in a single probe trial at 10 days post-learning (**Figure 6g**). These short term and long term neuroinflammatory induced memory deficits were both reversed by i.p. treatment by delivery to MG/Mφ of the Wnt antagonist, L803 mts (3DNA L803 mts Cy3) by the 3DNA nanocarrier (**Figure 6f,g**).

Altogether these data clearly demonstrate that a Wnt agonist therapy can reduce pro-inflammatory MG/Mφ activation and improve neuropathological and functional outcomes in a model of neuroinflammation mediated white matter injury. These data also highlight a novel 3DNA nanocarrier that has the ability to deliver therapies like this *in vivo* specifically to MG.

## Discussion

Recent research has highlighted that microglia of the developing brain have a unique phenotype. The aims of this study were, the identification in MG of the developing brain of the molecular regulators of injury-related phenotype, and validation of these as therapeutic targets for preventing microglia-mediated injury in the 15 million infants born preterm every year. We undertook a comprehensive set of studies involving pharmacologic and genetic manipulations in mouse and zebrafish models, human tissue and patient data. We also introduced an innovative 3DNA nanocarrier that can deliver drugs specifically to microglia *in vivo*. Initially, we characterised the complex MG/Mφ phenotype in our mouse model of encephalopathy of prematurity(Favrais et al 2011, Krishnan et al 2017, Schang et al 2014) and demonstrated that MG/Mφ activation is directly responsible in this model for the clinically relevant myelination defect. Next, using a genome-wide transcriptomic analysis of these activated MG/Mφ we associated the Wnt pathway with MG/Mφ activation. We specifically determined that the expression of Wnt/β-catenin pathway receptors, ligands and intracellular signalling components were robustly down-regulated in MG/Mφ isolated from our animal model. A combination of approaches in zebrafish and mouse models *in vitro* and *in vivo* allowed us to determine that Wnt/β-catenin pathway inhibition specific to MG/Mφ is sufficient and necessary to induce in these cells a pro-inflammatory activation, and subsequent hypomyelination *in vivo*. We highlighted the relevance of the *WNT* pathway to the activation state of human primary MG, and preterm born infant brain development. Of note, we verified our prediction that genetic variation in the *WNT* pathway in preterm infants is related to indices of their cerebral structural connectivity, although it is a limitation of these types of human studies that the data is not cell specific. Finally, we selectively delivered to MG/Mφ *in vivo* a Wnt pathway activator using a 3DNA nanocarrier, which we administered via non-invasive i.p. injection. This specific and non-invasive delivery of a Wnt agonist therapeutic prevented a pro-inflammatory MG/Mφ activation state and the associated white matter damage and cognitive deficit in our mouse model. Together, these data demonstrate that down-regulation of the canonical Wnt/β-catenin pathway triggers an oligodendrocyte-damaging MG phenotype, and validate this pathway as a novel and clinically viable neurotherapeutic target thanks to the MG/Mφ-specificity of 3DNA nanocarriers.

Our evidence supports that the Wnt pathway controlling MG/Mφ phenotype is the canonical Wnt/β-catenin pathway. Our supporting data include that genetic and pharmacological manipulation of the β-catenin degradation complex via Axin2 robustly effected MG/Mφ phenotype *in vitro* and *in vivo*. In contrast, blockade of non-canonical Wnt signalling with a PKC inhibitor had no effect on MG activation. Although, we acknowledge that there may still be a role for other facets of non-canonical signalling in the complexity of MG/Mφ functions. However, MG/Mφ-specific β-catenin ablation *in vivo* in-of-itself was sufficient to recapitulate the systemic-inflammation-driven hypomyelination that we see in our animal model of encephalopathy of prematurity. We also found that the WNT genes with variance associated with preterm infant connectivity were predominantly (8/10) from the Wnt/β-catenin pathway. A key role for β-catenin would also begin to explain our observations of Wnt-mediated pro-inflammatory activation via TLR-4 and IL-1 receptor agonists and also conversely Wnt-mediated anti-inflammatory activation via a IL-4 receptor agonist. Down-stream signalling of both the TLR/IL-1 and IL-4 receptor families includes activation of the transcription factor NF-κB (nuclear factor kappa-light-chain-enhancer of activated B cells)(Caamano & Hunter 2002). The ability of NF-kB to mediate the opposing effects of these receptors on MG phenotype is conferred by interactions with other regulatory factors, including at multiple levels indirectly and directly by β-catenin(Ma & Hottiger 2016).

Ours is the first study to demonstrate a robust cell-intrinsic role of the Wnt/β-catenin pathway in MG activation *in vitro* and *in vivo* and validate that this pathway is a viable immunomodulatory neurotherapeutic. Previous work on Wnt and MG has been limited to *in vitro* ligand-receptor interactions, specifically WNT3a or WNT5a. However, the specific Wnt pathway activated by a Wnt ligand is dependent on the context of ligand-receptor interaction not only cell-intrinsic properties(van Amerongen et al 2008). This fact likely explains the discordance between the data from the previous studies, where in some paradigms Wnt exposure is anti-inflammatory(Halleskog et al 2012, Yu et al 2014) or in others the exposure was pro-inflammatory(Halleskog et al 2011, Halleskog & Schulte 2013, Hooper et al 2012) with no relation to the predicted canonical or non-canonical pathway being activated. Altogether these data highlight the difficulty that would be faced in using ligand-receptor interactions to target the Wnt pathway to modulate MG activity, in contrast with our success of targeting the intracellular cascade of Wnt/β-catenin pathway specifically using our 3DNA-mediated delivery system.

The 3DNA nanocarrier is a powerful translational tool as is it can take agents across the BBB following i.p. injection, is specifically internalized by MG/Mφ in the brain and it has no toxicity *in vitro* or *in vivo* in our study, or in human retinal tissue *in vitro*(Gerhart et al 2017) and in pre-clinical *in vivo* targeted delivery of a ovarian cancer therapy(Huang et al 2016). Wnt activation in OPCs has previously been suggested to block their maturation by a member of our current research team(Fancy et al 2011). As such it was important that 3DNA-nanocarriers allowed us to target this pathway exclusively in MG/Mφ. 3DNA has a non-injury dependent mechanism of traversing the BBB, as it crossed in naïve and inflammation-exposed mice, although the specific mechanism is unknown. We have previously reported that the BBB in our mouse model of encephalopathy of prematurity is primarily intact(Krishnan et al 2017). The chemical and physical attributes of 3DNA likely precludes it from entering the brain via paracellular aqueous routes or transcellular lipophilic pathways. However, investigations of receptor mediated transcytosis are underway, as similar aptamer-based DNA duplexes can interact with receptors such as nucleolin or transferrin(Reyes-Reyes et al 2010).

In preterm infants, it is exposure to difficult to identify or prevent inflammatory events, such as chorioamnionitis and sepsis, that is the leading risk factor for white matter damage. This damage is caused by microglia-mediated neuroinflammatory processes(Hagberg et al 2015). White matter damage in these infants is observed as reduced structural connectivity on MRI(Shah et al 2008). This reduced structural connectivity is in turn strongly associated with poor neurodevelopmental outcomes(Ball et al 2015, Woodward et al 2006). As such, the purpose of our imaging-genomics analysis was to link these facts and to determine if genetic variance that could alter the microglial inflammatory response in our preterm cohort would be linked to changes in structural connectivity. Our integrated analysis of imaging and genomics demonstrated that common genetic variation in *WNT* pathway genes does indeed influence brain structural connectivity features within our preterm born infants. We fully acknowledge that the data we derived is not cell specific. However, it supports in contemporaneous infants the relevance of the *WNT* pathway using an imaging modality that is currently used to assess injury and predict outcome. In addition, we found that the *WNT* pathway genes containing SNPs associated with white matter tractography phenotype belonged to a human brain-specific gene interaction network. This network further builds a case for a functional effect of *WNT* SNPs in human preterm infants, with consequences for the development of white matter structure(Greene et al 2015). Specific prediction of the consequences of our identified SNPs found the majority were intron variants. Previous studies in experimental animals has shown that immune cell intron variants often effect cell function by altering enhancer binding(Farh et al 2015). Altogether our clinical data build a case that these *WNT* pathway SNPs may be a useful way to stratify infants who are at highest risk for white matter damage, and to improve on our currently limited prognostic abilities related to long term outcome.

In conclusion, the activity of the Wnt/β-catenin pathway regulates MG/Mφ activation, the Wnt pathway is relevant in human brain development, and specific agonism of the Wnt/β-catenin pathway in MG/Mφ *in vivo* using 3DNA nanocarrier-mediated drug-delivery has a beneficial effect on MG/Mφ phenotype, myelination and cognitive deficits. This is strong evidence that Wnt signalling modulation may be a promising therapeutic approach in encephalopathy of prematurity and, potentially, in other disorders involving MG/Mφ-mediated neuroinflammation.

## Methods

### Study approval

Experimental protocols were approved by the institutional guidelines of the Institut National de la Santé et de la Recherché Scientifique (Inserm, France) (2012-15/676-0079), and met the guidelines for the United States Public Health Service’s Policy on Humane Care and Use of Laboratory Animals (NIH, Bethesda, Maryland, USA).

The EPRIME study was reviewed and approved by the National Research Ethics Service, and all infants were studied following written consent from their parents.

For human MG cell culture, all procedures had ethical approval from Agence de Biomédicine (Approval PFS12-0011). Written informed consent was received from the tissue donor.

### Nomenclature of MG phenotype

We have adopted nomenclature consistent with our previous work in primary MG(Chhor et al 2013) acknowledging that this is a simplification necessary to facilitate description of the data. To simplify, we distinguished 3 types of phenotypes according to the mRNA expression levels of markers listed in brackets: Pro-inflammatory (*Nos2, Ptgs2, Cd32, Cd86* and *Tnf*α), anti-inflammatory (*Arg1, Cd206, Lgals3, Igf1, Il4 and Il10*) and immunoregulatory (*Il1rn, Il4ra, Socs3* and *Sphk1*).

### Mice

Experiments were performed with male OF1 strain mice pups from adult females purchased from Charles River (L’Arbresle, France) or Transgenic (Tg) male mice born in our animal facility. See the justification below in animal model section regarding use of male mice only. Tg mice B6.129P-Cx3cr1^tm^1^Litt^/J (CX3CR1-GFP) mice, B6.129-Ctnnb1^tm2Kem^/KnwJ (β-catenin^lfox^) mice and B6.129P2-Lyz2^tm^1^(cre)Ifo^/J (LysM^Cre^) mice were purchased from The Jackson Laboratory (ME, USA). CX3CR1-GFP mice express EGFP in monocytes, dendritic cells, NK cells, and brain MG under control of the endogenous Cx3cr1 locus. Only heterozygous (Cx3cr1^GFP/+^ and β-catenin^Δ/+^) mice were used in this study. Cx3cr1^GFP/+^ are obtained by crossing Cx3cr1^GFP/GFP^ with C57bl6J (WT) mice. LysM^Cre^ mice are useful for Cre-lox studies of the myeloid cell lineage. Here, we used it to analyse β-catenin deficit in the brain MG by crossing β-catenin^flox^ with LysM^Cre^ mice. Control mice were obtained by crossing β-catenin^flox^ with C57bl6J (WT) mice.

### Transgenic fish lines and fish maintenance

Tg (pu1::Gal4-UAS-TagRFP)(Sieger et al 2012) fish were kept at 26.5 °C in a 14 hour light/10 hour dark cycle. Embryos were raised at 28.5 °C in E3-solution until 120 hpf for analyses.

### Drugs

Mouse cytokines (IL-1β, IL-4) were purchased from R&D systems (France). LPS-EB Ultrapure was purchased from InvivoGen (France). To inhibit the Wnt/β-catenin pathway, we used XAV939 (Sigma-Aldrich, France). To activate it, we used CT99021 (Selleckchem, TX, USA), Lithium Chloride (Sigma-Aldrich, France), L803mts (Tocris, UK). The L803mts peptide and a scrambled peptide (SCR) were purchased conjugated to short oligonucleotide with disulphide bond linker from BioSynthesis (TX, USA) then hybridized to Cy3-labelled 3DNA at Genisphere (PA, USA). Chelerythrine was purchased from Sigma-Aldrich.

### Model of Encephalopathy of prematurity induced by IL-1β

Mice were housed under a 12-hour light-dark cycle, had access to food and water ad libitum and were weaned into same sex groups at P21. On P1 pups were sexed and where necessary litters were culled to 6-8 (Tg mice) or 12 (OF1 mice) pups. Assessments of injury and outcomes were made only in male animals and all pups within a litter received identical treatment to reduce any effects of differing maternal care. IL-1β exposure was carried out as previously described(Favrais et al 2011). Male mice only are used, as only male pups display a myelin deficit following exposure to IL-1β in this paradigm. This higher severity of injury in male animals is in accordance with observations of a greater burden of neurological injury in male infants following perinatal insult (Johnston & Hagberg 2007). A 5 µl volume of phosphate-buffered saline (PBS) containing 10 µg/kg/injection of recombinant mouse IL-1β (R&D Systems, MN, USA) or of PBS alone (control) was injected intra-peritoneally (i.p.) in male pups twice a day (morning and evening) on days P1 to P4 and once in the morning at P5.

### Morphological analysis of MG

P1 and P3 brain from Cx3Cr1-GFP mice administered PBS or IL-1β were fixed for 24 hours in 4% buffered formalin (QPath, Labonord SAS, France). Following fixation, cerebral tissue was sliced along a sagittal plane on a calibrated vibratome (VT1000 S, Leica, Germany) into 100 μm thick free-floating slices. Morphological analysis of MG was assessed as previously described(Verdonk et al 2016), using spinning disk confocal system (Cell Voyager - CV1000, Japan) with a UPLSAPO 40×;/NA 0.9 objective and the use of a 488-nm laser. A mosaic of more than one hundred pictures covering approximately 3.17 mm^2^ of tissue (cortex and white matter) per brain. An automatic image analysis using a custom designed script developed with the Acapella™ image analysis software (version 2.7 - Perkin Elmer Technologies) was realize to obtain density, cell body area, number of primary, secondary and tertiary processes and the area covered by the MG processes in 2D. A complexity index was calculated to analyse the morphological modifications induced by IL-1β.

*Neural tissue dissociation and CD11B+ MG or O4+ oligodendrocytes magnetic-activated cell sorting* At P1, P2, P3, P5 and P10, brains were collected for cell dissociation and CD11B+ cell separation using a magnetic coupled antibody anti-CD11B (MACS Technology), according to the manufacturer’s protocol (Miltenyi Biotec, Germany) and as previously described(Schang et al 2014). In brief, mice were intracardially perfused with NaCl 0,9%. After removing the cerebellum and olfactory bulbs, the brains were pooled (n=2-3 per sample) and dissociated using the Neural Tissue Dissociation Kit containing papain and the gentleMACS Octo Dissociator with Heaters. MG were enriched using the anti-CD11B (MG) MicroBeads, pelleted and conserved at −80 °C. The purity of the eluted CD11B+ fraction was verified as previously described (*15*). At P5 and P10, brains were collected for, O4+ oligodendrocytes cell separation using a magnetic coupled antibody anti-O4 (MACS Technology), according to the manufacturer’s protocol (Miltenyi Biotec, Germany) and as previously described (Schang et al 2014).

### RNA extraction and quantification of gene expression by real-time qPCR

Total RNA was extracted with the RNeasy mini kit according to the manufacturer’s instructions (Qiagen, France). RNA quality and concentration were assessed by spectrophotometry with the NanodropTM apparatus (Thermofisher Scientific, MA, USA). Total RNA (500ng) was subjected to reverse transcription using the iScriptTM cDNA synthesis kit (Bio-Rad, France). RT-qPCR was performed in triplicate for each sample using SYBR Green Super-mix (Bio-Rad) for 40 cycles with a 2-step program (5 s of denaturation at 95°C and 10 s of annealing at 60°C). Amplification specificity was assessed with a melting curve analysis. Primers were designed using Primer3 plus software (See sequences in **Supp Table 8)**. Specific mRNA levels were calculated after normalization of the results for each sample with those for Rpl13a mRNA (reference gene). The data are presented as relative mRNA units with respect to control group (expressed as fold over control value).

### Immunohistofluorescence

For frozen sections, P5 mice were intracardially perfused with 4% paraformaldehyde in 0.12M phosphate buffer solution under isoflurane anaesthesia. Brains were then post-fixed in 4% paraformaldehyde (Sigma-Aldrich, France) overnight at 4°C After 2 days in 30% sucrose 0.12M phosphate buffer solution, brains were frozen at −45°C in isopentane (Sigma-Aldrich) before storage at −80°C until sectioning on a cryostat (thickness14μM). Immunohistofluorescence was performed as previously described(Miron et al 2013). Slides were permeabilized and blocked for 1 hour and primary antibody was applied overnight at 4 °C in a humid chamber. To detect MG, rabbit antibody to IBA1 (Wako, 1: 500) was used. To detect proinflammatory MG we used mouse antibody to iNOS (BD Biosciences, 610329, 1:100), rat antibody to CD16/CD32 (BD Pharmingen, 553141, 1:500), rabbit antibody to COX2 (Abcam, Ab15191, 1:500). To detect anti-inflammatory MG, we used goat antibody to Arginase-1 (ARG1) (Santa Cruz Biotechnology, sc-18355, 1:50), rabbit antibody to mannose receptor (MR) (Abcam, ab64693, 1:600). To detect MG and Mφ rat antibody to CD68 (Abcam, ab53444, 1:100) was used. Fluorescently conjugated secondary antibody to goat IgG (A21432), antibodies to rabbit IgG (A11034, A10042), antibodies to rat IgG (A21434, A11006, A21247) and antibody to mouse IgG (A21042), were applied for 2 hours at 20–25 °C in a humid chamber (1:500, Invitrogen). Following counterstaining with Hoechst, slides were coverslipped with Fluoromount-G (Southern Biotech, Cliniscience, France). Antibody isotype controls (Sigma-Aldrich) added to sections at the same final concentration as the respective primary antibodies showed little or no nonspecific staining. All manual cell counts were performed in a blinded manner.

### In vivo treatments with XAV939, 3DNA L803mts Cy3, and Gadolinium

XAV939 (250µM, 0.5nmol/injection) or PBS/DMSO (vehicle) and 3DNA L803mts Cy3 (200ng/injection) or 3DNA SCR Cy3 (200ng/injection) (control peptide) were injected into the right ventricle of mice pups at P1 using 26-G needle linked to a 10µL Hamilton syringe mounted on a micromanipulator and coupled to a micro-injector (Harvard Apparatus, MA, USA; outflow 2 µL/min), 3DNA SCR or L803 mts Cy3 were injected 1 hour before i.p. injection of PBS or IL-1β. Three hours after PBS or IL-1β injection or 4 hours after XAV939 or vehicle injection, pups were sacrificed for CD11B-postive cell sorting. Chronic treatment with 3DNA L803mts Cy3 were performed by i.p. co-injections of PBS or IL-1β with 3DNA L803mts Cy3 (500ng/injection) or 3DNA L803mts Cy3 (500ng/injection) twice a day between P1 and P3 and once a day at P4 and P5. Effect of this treatment on myelination was analysed at P10 and behavioural tests were realized in adulthood (2-3months). To analyse uptake of 3DNA in liver and spleen, one i.p. injection of 3DNA Cy3 (500ng/injection) with PBS or IL-1β was performed and animals sacrificed 4 hours later. To analyse uptake of 3DNA in brain, animals was sacrificed 4 hours after i.c.v. injection at P1, and at the end of the i.p. treatment at P5.

Selective depletion of pro-inflammatory MG in white matter injury was performed using gadolinium chloride (GdCl3) as previously described(Miron et al 2013) with slight modifications. Briefly, PBS or Gadolinium (200nmol/injection, Sigma, France) was injected into the corpus callosum of mice pups at P1, 1 hour before i.p. injections of PBS or IL-1 β using 26-G needle linked to a 10µL Hamilton syringe mounted on a micromanipulator and coupled to a micro-injector (Harvard apparatus, outflow 2 µL/min). Effect of pro-inflammatory MG depletion on MG cell phenotype and myelination was analysed at P10.

### Actimetry

The horizontal (spontaneous locomotion) and vertical (rearing) activities were individually assessed in transparent activity cages (20×10×12cm) with automatic monitoring of photocell beam breaks (Imetronic, Bordeaux, France). Actimetry test was performed at ≈ 2 months (day 65), with recording every 30 minutes over 23 hours, in order to evaluate the effect of IL-1β and 3DNA treatments i.p. injections on spontaneous locomotor activity, as modified locomotor activity could lead to biased results from other behavioural tests requiring locomotion.

### Open-field

The open-field test was performed at ≈ 2.5 months (day 80) using a square white open-field apparatus (100×100×50cm) made of plastic permeable to infrared light. Distance travelled and time spent in inactivity were recorded by a videotrack system (Viewpoint^®^) during the 9 minutes’ test.

### Barnes maze test

The Barnes maze test was performed at 3 months (day 90 to 94). The maze is a wet, white and circular platform (80cm diameter), brightly illuminated (400lux), raised 50cm above the floor, with 18 holes (4.5cm) equally spaced around the perimeter. A white hidden escape box (4×13×7cm), representing the target, was located under one of the holes. Prior to the test, each mouse was subjected to a habituation trial where the mouse was directly put in the escape box for 30 seconds. Learning: the mouse was placed in the centre of the circular maze and was allowed to explore the platform and holes for 3 minutes’ maximum. The distance travelled was recorded using a videotrack system (Viewpoint^®^). Latency to reach the escape box, and number of errors (number of empty holes visited) were manually noted. When the mouse found and entered the escape box, the videotrack recording was stopped, and the mouse remained in the escape box for an additional 10 seconds before returning to its home cage. If the mouse did not enter spontaneously, it is gently put toward the escape box, before returning to its home cage. On the first day of training, mice underwent 2 trials after the habituation trial; thereafter, 3 trials were given per day, with a 2 hours’ interval.

Probe trial: on the 5th day of learning, the probe trial, with the escape box removed and lasting 90 seconds, is used to assess spatial memory performances. Time spent and distance travelled in each sextant (defined by one of the 6 parts of the maze, that includes 3 holes) were recorded. The target zone is defined as the part, which contains the target hole and two adjacent holes.

Long term trial: 15 days after learning, a new trial, with the escape box, is used to assess long-term retention memory. Escape latency and distance travelled to reach the target were recorded.

### Immunohistochemistry

Brains were collected at P15 or P30 and immersed immediately after sacrifice in 4% formaldehyde for 4 days at room temperature, prior to dehydration and paraffin embedding. Section was realized using a microtome. Immunostaining was performed as previously described(Favrais et al 2011) using mouse antibody to Myelin Basic Protein (MAB382, Millipore, France). The intensity of the MBP immunostaining in corpus callosum was assessed by a densitometry analysis through NIH ImageJ Software (http://imagej.nih.gov/ij/). Optical density was deduced from grey scale standardized to the photomicrograph background. Corpus callosum was defined as region of interest. One measurement per section and 4 sections were analysed in each brain.

### Primary mouse MG culture

Primary mixed glial cell cultures were prepared from the cortices of postnatal day 1 OF1 mice, as previously described(Chhor et al 2013). After 14 days, MG were purified and pelleted via centrifugation and re-suspended in DMEM/PS/10% FBS at a concentration of 4×10^5^ cells/mL. One or two mL/well of cell suspension was plated in 12-well (for qRT-PCR) or 6-well (for ELISA) culture plates respectively. For immunofluorescence, 250 µL of cell suspension was plated in μ-Slide 8 Well Glass Bottom (Ibidi, Biovalley, France). Post-plating, media was replaced after 1 hour. Based on previous work from our lab(Chhor et al 2013), the day after plating, MG were exposed for 4 hours to DMEM (control) or IL-1β at 50 ng/mL, IL-4 at 20 ng/mL, diluted in DMEM. For RT-qPCR or ELISA analysis, media were removed and plates frozen at −80°C. For immunofluorescence experiments, cells were fixed at room temperature with 4% paraformaldehyde for 20 minutes.

### Mixed mouse glial and OPC cultures

Cortical mixed glial cultures and purified OPC were obtained from the cortices of postnatal day 1 OF1 mice as previously described with slight modification (*40*). Mixed glial cells were resuspended in MEM-C, which consists of Minimal Essential Medium (MEM) supplemented with 10% FBS (Gibco), Glutamax (Gibco), and 1% PS, and plated at 2×10^5^ cells/cm^2^ onto T75 flasks for subsequent OPC purification or Ibidi 8-well chamber slides (Biovalley, France) for mixed glial culture treatment. For this purpose, mixed glial cultures were maintained in proliferation in MEM-C for 7 days, then maintained during 10-12 days in differentiation medium which consists of Dulbecco’s Modified Eagle Medium (DMEM-F12) with glutamine (Gibco), N2 (Gibco) and 1% PS. Half of the wells were exposed to IL-1β (50 ng/ml) and medium was changed every 2-3 days. Alternatively, mixed glial cultures plated onto T75 flasks were maintained for 9-12 days in MEM-C and purified OPC cultures were prepared by a differential shake(McCarthy & de Vellis 1980). OPCs were seeded onto Ibidi 8-well chamber slides at a density of 3 ×; 10^4^ cells/cm^2^ in proliferation medium which consists of MACS Neuro Medium with Neurobrew 21 (Miltenyi Biotec), Glutamax, 1% PS, FGFb and PDGFa (10 ng/ml each, Sigma, France). After 72h, differentiation was induced by FGFb and PDGFa withdrawal and by adding T3 (40 ng/ml, Sigma). Half of the wells were exposed to IL-1β (50 ng/ml). After 10-12 days (for mixed glial cultures) or 72h (for OPCs) of IL-1β, cells were fixed 20 min with 4% paraformaldehyde.

### Primary human MG culture

Within 1 hour of scheduled termination (age 19 and 21 weeks of amenorrhea) brain tissues were collected from two human foetuses without any neuropathological alterations. Brain tissue (4 g) was minced and further mechanically dissociated using 1ml micropipettor in HBSS with Ca^2+^ and Mg^2+.^ Using a 70µM strainer a single cell suspension was obtained. MG/Mφ isolation were obtained using anti-CD11B (MG) microbeads (MACS Technology), according to the manufacturer’s protocol (Miltenyi Biotec, Germany), as above. CD11B+ MG/Mφ were pelleted via centrifugation and re-suspended in DMEM/PS/10% FBS at a concentration of 5×10^6^ cells/mL. Cells were plated in 12-well plates (1ml/well) for qRT-PCR analysis. For immunofluorescence analysis, cells were plated in μ-Slide 8 Well Glass Bottom (Ibidi, Biovalley, France). 48 hours after plating, MG were exposed for 4 hours to DMEM (control) or LPS 10 ng/mL diluted in DMEM. For RT-qPCR media were removed and plates frozen at −80°C. For immunofluorescence experiments, cells were fixed at room temperature with 4% paraformaldehyde for 20 minutes.

### Immunocytofluorescence

Immunofluorescence staining was performed as previously described(Favrais et al 2011), using mouse antibody to MBP (MAB382, 1: 500; Millipore), rabbit antibody to OLIG2 (18953, 1:500 IBL-America, MN, USA), rabbit antibody to beta-catenin (E247, 1:100; Millipore), rabbit antibody to COX2 (Ab15191, 1: 400; Abcam, MA, USA) goat antibody to ARG1 (sc 18354, 1:400; Santa-Cruz, CA, USA) and IL1RA (sc 8482, 1:200, Santa-Cruz). Fluorescently conjugated secondary antibody (all Thermfisher Scientific) to mouse IgG (A21236), to rabbit IgG (A21206) and to goat IgG (A11055), were applied for 2 hours at 20–25 °C in a humid chamber (1:500, Invitrogen). Slides were coverslipped with Fluoromount-G (Southern Biotech, AL, USA). To analyse MBP labelling, MBP area was assessed as the percentage of total area above threshold using NIH ImageJ software (http://imagej.nih.gov/ij/) and was normalized to the number of Olig2-positive cells. Means were expressed as fold change over control condition.

### Microarrays and data pre-processing

Gene expression quantification, including RNA extraction and quality assurance, was performed by Miltenyi Biotec on a total of 24 samples of CD11B+ cells and 20 samples of O4+ cells from the brains of mice exposed to IL-1β or PBS between P1 and P5, and sacrificed at P5 or P10. RNA was extracted and hybridised to Agilent Whole Mouse Genome Oligo Microarrays (8×60K).

### Bioinformatics’ gene co-expression network reconstruction

#### Expression data

Each cell type and experimental condition was analysed separately, to investigate changes in gene expression over time in response to exposure to IL-1β or PBS. Expression values for MG (CD11B+) and oligodendrocytes (O4+) at P5 and P10 were quantile normalized.

#### Differential expression

Probes showing high variability across time points within each cell type were retained for further analysis (coefficient of variation above 50th centile). Probes showing differential expression between time points were identified using unpaired t-tests with Benjamini-Hochberg correction for multiple testing in the Bioconductor multitest package(Opgen-Rhein & Strimmer 2007, Schafer & Strimmer 2005)

(Gentleman et al 2004), with a false discovery rate threshold (FDR) of 10%.

#### Network reconstruction

The co-expression network was inferred using Graphical Gaussian Models (GGM), implemented within the software package “GeneNet”(Schafer & Strimmer 2005) in R (http://www.r-project.org/). This analysis computes partial correlations, which are a measure of conditional independence between two genes i.e. the correlation between the expression profiles of two genes after the common effects of all other genes are removed. Correction was made for multiple comparisons by setting the local false discovery rate (FDR) at ≤1%. Individual networks were reconstructed for temporally differentially expressed genes for each cell type in each experimental condition (i.e. 2 cell types x 2 conditions). The input for each GGM was a matrix of differentially expressed normalised mRNA transcript levels. The genes in each of these four co-expression networks (**Supp Figure 3e**) were functionally annotated by calculating significant enrichment of KEGG pathways using the Broad Institute MSigDB database (http://software.broadinstitute.org/gsea/msigdb/index.jsp)(Subramanian et al 2005) (**Supp Table 2)**.

Differential expression analysis was performed using the limma package (Smyth 2004). The mouse datasets were annotated with human Ensembl gene ID using the biomaRt Bioconductor R package(Durinck et al 2009) and selecting human genes that were ‘one-to-one’ orthologues with mouse genes. We multiplied the adjusted P value (the –log10 of P value) by the log-transformed fold change to generate a gene-level score, which was used as a metric to ‘rank’ genes. GSEA (Subramanian et al 2005) was applied genome-wide to the ranked list of gene scores (reflecting both the significance and the magnitude of expression changes in IL-1β exposed MG/Mφ) to test if a group of genes (MSigDB gene sets) occupy higher (or lower) positions in the ranked gene list than what it would be expected by chance. Gene set enrichment scores and significance level of the enrichment (NES, P value, FDR) and enrichment plots were provided in the GSEA output format developed by Broad Institute of MIT and Harvard (permutations = 10,000). The GSEA was carried out accounting for the direction of IL-1β effect, i.e. considering whether a gene is upregulated or down-regulated under IL-1β exposure at P5 and P10.

### Wnt/β-catenin pathway modulation in primary MG

MG transfection was realized using negative control siRNA or siRNA Axin2 (ON TARGET Plus Control Pool and ON TARGET Plus mouse Axin2-06, Thermoscientific, Dharmacon, France) at a final concentration of 30µM. Transfection was realized using the MACSfectin transfection reagent (Miltenyi biotec, Germany) in mixture of Opti-MEM and DMEM mediums for 48 hours prior to IL-1β stimulation. *Axin2* mRNA knockdown was evaluated by qRT-PCR. XAV939 and CT99021 (1µM) or PBS/DMSO (vehicle) were added to media 30 minutes before DMEM (control) or IL-1β (50 ng/ml) treatments.

### PSer45 β-Catenin and β-Catenin quantification by ELISA

Quantification of PSer45 β-Catenin and β-Catenin by ELISA was realized using β-catenin pSer45 + Total PhosphoTracer ELISA Kit (ab119656, Abcam) for primary microglia or β-catenin ELISA Kit (ab1205704, Abcam) for MACSed cells. Briefly, cell lysates were obtained using lysis buffer of the kit. Total protein level was quantified using BCA method (Bicinchoninic Acid Solution and Copper (II) Sulfate Solution from Sigma, France). The ELISA was realized according to the manufacturer’s instruction. Quantification of PSer45 β-Catenin is a ratio PSer45 β-Catenin/total β-Catenin. Total *β*-Catenin was normalized using total protein level. Total β-Catenin data are expressed as fold over control values.

### Wnt/β-catenin pathway modulation in Zebrafish

MG were visualized using pu1::Gal4-UAS::TagRFP. 72 hpf old zebrafish embryos were used for LPS microinjection (5ng/injection) in hindbrain. The Wnt/β-catenin pathway was modulated by adding LiCl (80mM), XAV939 (5μM), CT99021 (3μM) in E3-solution. Fluorescently labelled embryos were imaged using a microscope equipped with an ApoTome system (Zeiss, France) fitted with an AxioCam MRm camera (Zeiss) controlled by the Axiovision software. The thickness of the z stacks was always comprised between 2.5 and 3 μm. Fluorescence intensity was quantified by NIH ImageJ software (http://imagej.nih.gov/ij/) on grey scale images.

### Preterm infant cohort

Preterm infants were recruited as part of the EPRIME (Evaluation of Magnetic Resonance (MR) Imaging to Predict Neurodevelopmental Impairment in Preterm Infants) study and were imaged at term equivalent age over a 3-year period (2010-2013) at a single centre. The EPRIME study was reviewed and approved by the National Research Ethics Service, and all infants were studied following written consent from their parents. 290 infants (gestational age 23.57 to 32.86 weeks, median 30 weeks) were scanned at term-equivalent age (38.29 to 58.28 weeks). Pulse oximetry, temperature, and heart rate were monitored throughout the period of image acquisition; ear protection in the form of silicone-based putty placed in the external ear (President Putty, Coltene, UK) and Mini-muffs (Natus Medical Inc., CA, USA) were used for each infant.

### Image Acquisition

MRI was performed on a Philips 3-Tesla system (Best, Netherlands) using an 8-channel phased array head coil. The 3D-MPRAGE and high-resolution T2-weighted fast spin echo images were obtained before diffusion tensor imaging. Single-shot EPI DTI was acquired in the transverse plane in 32 non-collinear directions using the following parameters: repetition time (TR): 8000 ms; echo time (TE): 49 ms; slice thickness: 2 mm; field of view: 224 mm; matrix: 128 ×; 128 (voxel size: 1.75 ×; 1.75 ×; 2 mm^3^); *b* value: 750 s/mm^2^. Data were acquired with a SENSE factor of 2.

### Data selection and quality control

The T2-weighted MRI anatomical scans were reviewed in order to exclude subjects with extensive brain abnormalities, major focal destructive parenchymal lesions, multiple punctate white matter lesions or white matter cysts. All MR-images were assessed for the presence of image artefact (inferior-temporal signal dropout, aliasing, field inhomogeneity, etc.) and severe motion (for head-motion criteria see below). 290 infants had images suitable for tractography and associated genetic information.

DTI analysis was performed using FMRIB’s Diffusion Toolbox (FDT v2.0). Images were registered to their non-diffusion weighted (*b*0) image and corrected for differences in spatial distortion due to eddy currents. Non-brain tissue was removed using the brain extraction tool (BET)(Smith 2002). Diffusion tensors were calculated voxel-wise, using a simple least-squares fit of the tensor model to the diffusion data. From this, the tensor eigenvalues and FA maps were calculated.

### Probabilistic tractography

Regions of interest for seeding tracts were obtained by segmentation of the brain based on a 90-node anatomical neonatal atlas(Shi et al 2011), and the resulting segmentations were registered to the diffusion space using a custom neonatal pipeline(Ball et al 2010). Tractography was performed on DTI data using a modified probabilistic tractography which estimates diffusive transfer between voxels(Robinson et al 2008), using cortico-cortical connections only. A weighted adjacency matrix of brain regions was produced for each infant: self-connections along the diagonal were removed; symmetry was enforced; and the redundant lower triangle removed. The edges of these connectivity graphs were vectorised by concatenating the rows for each individual connectivity matrix, and appending them to form a single matrix of *n* individuals by *p* edges. Each phenotype matrix was adjusted for major covariates (post-menstrual age at scan and at birth) and reduced to its first principal component.

### Genome-wide genotyping

Saliva samples from 290 preterm infants with imaging data were collected using Oragene DNA OG-250 kits (DNAGenotek Inc., Kanata, Canada) and genotyped on Illumina HumanOmniExpress-24 v1.1 chip (Illumina, San Diego, CA, USA). Filtering was carried out using PLINK (Software: https://www.cog-genomics.org/plink2). All individuals had missing call frequency < 0.1. SNPs with Hardy-Weinberg equilibrium exact test p≥1 ×; 10-6, MAF ≥ 0.01 and genotyping rate > 0.99 were retained for analysis. After these filtering steps, 659 674 SNPs remained.

### Preterm infant imaging genomics analysis of WNT pathway

A list of all genes in the *WNT* signalling pathway (entry hsa04310) in the Kyoto Encyclopaedia of Genes and Genomes (KEGG) was used as a gene-set-of-interest (*n*=141 genes). Gene-set analysis was carried out with the Joint Association of Genetic Variants (JAG) tool(Lips et al 2012, Lips et al 2015) to test the joint effect of all SNPs located in the WNT pathway. This procedure consists of three parts: 1) SNP to gene annotation; 2) self-contained testing (i.e. association) and 3) competitive testing (i.e. enrichment). An enriched pathway can be defined as one whose genes are more strongly associated with the phenotype than those genes outside the pathway. Phenotype permutation is used to evaluate gene-set significance, implicitly controlling for linkage disequilibrium, sample size, gene size, number of SNPs per gene, and number of genes in a gene-set. This will result in a *p*-value (with a traditional significance threshold<0.05) if there is an enrichment for variants associated with the phenotype within the gene-set of interest compared to random matched gene-sets.

Following SNP-gene mapping, genes in the WNT-set were tested for association with the original phenotype (query data), and this was repeated with 1000 permutations of the phenotype (self-contained/association test). This tests the null hypothesis that no pathway genes are associated with the phenotype, and combines the individual gene association p-values into a single p-value for the entire pathway. The competitive/enrichment test was then done to test the null hypothesis that the pathway genes are no more associated with the phenotype than 300 randomly drawn non-pathway gene-sets, with 1000 phenotype permutations. The association analysis was repeated gene-by-gene within the *WNT* gene-set to investigate which members of the gene-set might be predominantly contributing to the collective signal.

### Statistical analysis

No statistical methods were used to predetermine sample sizes but these were similar to those generally employed in the field. Data are expressed as the mean values +/− standard error of the mean (SEM). Data was first tested for normality using the Kolmogorov-Smirnov normality test for n=5-7 and D’Agostino & Pearson omnibus normality test for n>7. F-test (Single comparisons) or Bartlett’s test (Multiple comparison) for analyse of variances was used. Multiple comparisons in the same data set were analysed by one-way ANOVA with Newman-Keuls *post hoc* test, Two-way ANOVA with Bonferroni *post hoc* or Kruskal-Wallis test with Dunns post hoc. Single comparisons to control were made using two-tailed Student’s *t*-test or Mann-Whitney test. *P* < 0.05 was considered to be statistically significant. For the Barnes maze probe trial, univariate t test was performed to compare the % time spent in the target sextant to the theoretical value 16.67 % (i.e. when the mouse spent equal time within each sextant). Data handling and statistical processing was performed using Microsoft Excel and GraphPad Prism Software.

## Author contributions

J.V.S., A.L.S., M.K., F.V., V.D., O.H., S.L., L.S., T.L.C., G.B., P.A., A.S., C.A., V.M.; D.R., F.C., C.L., V.B., E.G.P., A.D.E., H.H., N.S.Y., B.F., P.G. designed the research; J.V.S., A.L.S., M.L. K., S.S., A.M., F.V., O.H., C.B., A.D., S.L., L.S., T.L.C., R.H.A., G.B., P.A., A.S., R.K.H., C.A., V.M., C.L., V.B., N.S.Y., B.F., P.G. performed the research; W.B. provided the β-catenin floxed mice; J.V.S., A.L.S., M.L.K., S.S., A.M., F.V., V.D., O.H., S.L., L.S., T.L.C., R.H.A., G.B., P.A., A.S., C.A., V.M., C.B., A.D., D.R., F.C., C.L., V.B., E.G.P., A.D.E., H.H.H, N.S.Y., B.F., P.G. analysed the data; J.V.S., M.L.K., D.R., FC., C.B., A.D., C.L., V.B., E.G.P., A.D.E., H.H, N.S.Y., B.F., P.G. wrote the manuscript.

## Acknowledgments

This study was supported by grants from Inserm, Université Paris Diderot, Université Sorbonne-Paris-Cité, Investissement d’Avenir (ANR-11-INBS-0011, NeurATRIS), ERA-NET Neuron (Micromet), DHU PROTECT, Association Robert Debré, PremUP, Fondation de France, Fondation pour la Recherche sur le Cerveau, Fondation des Gueules Cassées, Roger de Spoelberch Foundation, Grace de Monaco Foundation, Leducq Foundation, Action Medical Research, Cerebral Palsy Alliance Research Foundation Australia, Wellcome Trust (WSCR P32674) and The Swedish Research Council (2015-02493). We wish to acknowledge the support of the Department of Perinatal Imaging & Health, King’s College London. In addition, the authors acknowledge financial support from the National Institute for Health Research (NIHR) Biomedical Research Centre based at Guy’s and St Thomas’ NHS Foundation Trust and King’s College London. The views expressed are those of the author(s) and not necessarily those of the NHS, the NIHR or the Department of Health

## Supplementary figures

**Supplementary Figure 1 related to Figure 2. Expression of phenotype markers by MG/M**φ **in IL-1**β **treated mice at P3. (a)** Quantification of CD16+, CD68+/iNOS+, CD68+/COX2+, CD68+/ARG1+ and MR (CD206)+ cells at P3 from PBS and IL-1β treated mice. Data are expressed as the number of positive cells per field and the number of animals is shown on the graph (Mann-Whitney test: mean±SEM; * p<0.05 and **p<0.01). Representative images at P3 in the corpus callosum of the effect of IL-1β on the number of **(b)** CD68+/iNOS+ and CD68+/ARG1+ cells **(c)** CD68+/COX2+ and **(d)** CD16+ and MR+ cells. Scale bar: 50μm

**Supplementary Figure 2 related to Figure 3 and Supp Table 1. Properties of the networks formed from the transcriptomic analysis of MG and oligodendrocytes following exposure to IL-1**β. In (**a**) the structural properties of the transcriptomic network from the microarray analysis performed on CD11B+ and O4+ cell fractions isolated by MACS. Reconstructed gene networks by cell type and condition, local FDR <1% (OG: oligodendrocytes; MG MG/Mφ; PBS: control). (**b**) Visual representation of the output of transcriptomic analysis of MG and oligodendrocytes following exposure to IL-1β. Under control conditions (PBS) the OG co-expression network appears less dense, structured with interconnected clusters (34 connected components) where most genes are not direct neighbours of one another (characteristic path length 5.5). With exposure to IL-1β there is a noticeable change in the OG network structure, so that there are many more genes being co-expressed (an increase from 571 nodes, 1229 edges in PBS to 1583 nodes, 6457 edges in IL-1β) and they are more highly interconnected (characteristic path length 2.9). The control (PBS) MG co-expression network is also relatively clustered and less dense, and on exposure to IL-1β there is a dramatic increase in co-expression (786 nodes, 1810 edges in PBS to 3113 nodes and 48104 edges in IL-1β). This demonstrates that at rest MG are more transcriptionally active than OG, and both cell-types respond strikingly to IL-1β exposure with a global activation of gene expression that is particularly pronounced for MG.

**Supplementary Figure 3 related to Figure 4. Validation in primary MG of an inverse relationship between pro-inflammatory status and the Wnt/**β**-catenin pathway.** In **(a)** MG phenotype induced by IL-1β was assessed via pro-inflammatory markers (*Cd32, Cd86, Nos2* and *Ptgs2*), anti-inflammatory markers (*Arg1, Cd206, Igf1* and *Lgals3*) and immuno-regulator markers (*Il1rn, Il4ra, Socs3* and *Sphk1*) by RT-qPCR after 4 hours of IL-1β stimulation. mRNA levels are presented as a fold change relative to PBS exposed MG. Data are expressed as mean±SEM, in brackets are for the n PBS and IL-1β groups (Student’s t-test: * p<0.05, ** p<0.01 and ***P<0.001). **(b)** MG phenotype induced after 4 hours of exposure to IL-1β in primary MG as assessed with immunoreactivity (IR) for COX2 (pro-inflammatory) ARG1 (anti-inflammatory) and IL1RA (immuno-regulator). Scale bar: 50μm. **(c)** Canonical Wnt pathway expression as altered by IL-1β exposure in primary MG as assessed by RT-qPCR for *Ctnnb1, Tcf1, Lef1* and Wnt receptors Frizzled (*Fzd*) *3,4,6* mRNA after 4 hours of stimulation. mRNA levels are presented as a fold change relative to control (PBS treated) MG. Data are expressed as the mean±SEM and n are indicated in the columns (Student’s t-test: * p<0.05, ** p<0.01 and *** p<0.001). **(d)** IL-1β induces down-regulation of β-catenin immunoreactivity in primary MG after 4 hours of stimulation. Scale bar: 50μm. **(e)** IL-1β induces down-regulation of total β-catenin without any modification of the PSer45 β-catenin/β-catenin ratio as assessed via ELISA (normalized to PBS control) in primary MG after 4 hours of stimulation. Data are expressed as the mean±SEM (Student t-test: * p<0.05, ** p<0.01 and *** p<0.001). **(f)** Expression of members of the canonical Wnt pathway in primary MG activated by IL-4 (20ng/ml) as assessed by RT-qPCR for *Ctnnb1, Tcf1* and *Lef1* after 4 hours of stimulation. mRNA levels are presented as a fold change relative to control (PBS treated) MG. Data are expressed as the mean±SEM, (Student‘s t-test: * p<0.05, ** p<0.01 and *** p<0.001). **(g)** IL-4 induces down-regulation of Pser45 β-catenin/β-catenin ratio without any modification of total β-catenin in primary MG as assessed with ELISA after 4 hours of stimulation. Data are expressed as the mean±SEM (Student’s t-test: * p<0.05, ** p<0.01 and*** p<0.001).

**Supplementary Figure 4. Non-canonical Wnt pathway blockade has no effect on the activation of primary MG stimulated by IL-1**β. Inhibition of Protein Kinase C by Chelerythrine has no effect on MG activation. Quantification of *Nos2, Ptgs2, Il1rn* and *Socs3* mRNA by RT-qPCR in control (PBS) and IL-1β treated primary MG with Chelerythrine (1 or 3µM) or DMSO. mRNA levels are presented as a fold change relative to control/DMSO group. (mean±SEM; # p<0.05, ## p<0.01, showed significant difference between PBS/DMSO and IL-1β/DMSO groups. Kruskal-Wallis test with Dunns *post-hoc* test). Data are from 3 independently replicated experiments.

**Supplementary Figure 5 related to Figure 6. Cell and organ specific analysis of the uptake by MG of 3DNA Cy3 labelled nanoparticles. (a)** 3DNA uptake by IBA-1+ cells in mixed primary cultures of astrocytes, OPCs and MG after incubation with 3DNA Cy3 (200ng/ml) for 24 hours. Scale bar: 40μm. **(b)** 3DNA uptake by a primary MG following incubation with 3DNA L803mts Cy3 (200ng/ml) for 5 hours. Scale bar: 10μm. Representative images of the undetectable uptake of 3DNA by the **(c)** spleen and **(d)** the liver. Scale bar: 100μm

**Supplementary Figure 6. Nanocarrier-mediated delivery of a Wnt/**β**-catenin pathway activator L803mts regulates MG/M**φ **activation in primary MG.** In (**a**) representative images of the *in vitro* uptake of 3DNA by primary MG 4 hours after 3DNA L803mts Cy3 exposure. In (**b**) effects of 3DNA L803mts Cy3 exposure to reduce the IL-1β-induced MG/Mφ activation as evaluated by qRT-PCR quantification of *Nos2, Ptgs2, Tnf*α, *Il1rn, Socs3* and *Il4ra* mRNA. mRNA levels are presented as a fold change relative to 3DNA SCR Cy3/PBS group (n for each data point is indicated on the bar, mean±SEM, One-way ANOVA with *post hoc* Newman-Keuls’s test: ++ p<0.01, +++p<0.001 indicate a statistically significant difference between the PBS and the IL-1β groups also exposed to 3DNA SCR Cy3. For the effects of Wnt modulator delivery, * p<0.05, ** p<0.01.

**Supplementary Figure 7. Nanocarrier-mediated delivery of a Wnt/**β**-catenin pathway activator L803mts has no effect on indices of basal behaviour in our encephalopathy of prematurity model.** In (**a**) Actimetry data including horizontal locomotion and rearing over a 24 hour period, mean±SEM. In (**b**) data on distance travelled and time spent inactive in the Open Field test, mean±SEM.

## Supplementary tables

**Supp Table 1.** Genes found within the co-expression networks for each of the four GGM outputs. See also, Supp Figure 1.

**Supp Table 2.** Biological pathway analysis using the Broad Institute MsigDB database for each of the four conditions. Of note for MG exposed to IL1B Wnt signalling is highly significantly enriched.

**Supp Table 3.** GSEA including genes down-reguated in the array analysis, showing a predominance of Wnt enrichment in the down-regulated genes.

**Supp Table 4.** Analysis of the 10 highly ranked genes showing statistical outputs in full for an association with the white matter connectivity phenotype.

**Supp Table 5.** Genes creating a coherent interaction network built around the 10 WNT pathway genes with SNPs highly associated with the preterm white matter connectivity phenotype.

**Supp Table 6.** Details for each of the 42 SNPs for analysis from the 10 genes most highly associated with the preterm white matter phenotype

**Supp Table 7.** Consequences predicted for the 42 SNPs found in the 10 genes of high relevance to the preterm white matter connectivity phenotype. Summary of effects found in Figure 5h.

**Supp Table 8.** List of primer sequences

